# Exposure to rotenone triggers redox driven system-wide lipidome alterations and metabolic trade-offs linked to Parkinson’s disease

**DOI:** 10.1101/2025.02.10.637453

**Authors:** Ashutosh Kumar Tiwari, Priya Rathor, Rajendra P Patel, Pawan Kumar Jha, Nick Birse, CH Ratnasekhar

## Abstract

With the global rise in aging populations, the increasing incidence of neurodegenerative diseases underscores concerns about brain health, with pesticides like rotenone, emerging as key environmental hazardous. The precise mechanism by which chronic environmental concentration of rotenone exposure causes Parkinson-like phenotype is not understood. Previous studies showed that rotenone induces depletion of dopaminergic neurons by influencing mitochondrial functions. Mitochondrial dysfunction alters lipid homeostasis; therefore, brain lipids can be potential targets for the early risk assessment and prognosis of Parkinson’s disease (PD). However, the specific lipidome changes and associated biomarkers of chronic rotenone in vivo exposure causing PD are largely unknown. This study investigates the lipid profile disruptions and biomarkers induced by environmentally relevant concentrations of chronic rotenone exposure using the neuro-model *Drosophila melanogaster*. An untargeted LC-HRAMS-based lipidomics identified that lipid classes, GP, SP, FA and GL were significantly altered. Furthermore, system-wide loss of cross talk of mitochondrial and peroxisome lipids by altering their redox homeostasis causing PD was observed. Additionally, lipid oxidative stress markers, and behavior abnormalities correlated with altered lipids linked to PD. The findings highlight the rotenone induced complex metabolic trade-offs, prioritizing brain’s neural integrity at the expense of peripheral lipid levels, leading to PD.

**Environmental implication:** This study highlights the environmental risks of chronic rotenone exposure, commonly used in agriculture. The findings show that even low concentrations of rotenone disrupt lipid metabolism, particularly in the brain, affecting mitochondrial and peroxisomal functions, contributing to depleting dopaminergic neurons, which are linked to neurodegenerative diseases like Parkinson’s. The study also identify lipid markers linked to rotenone induced PD and reveals metabolic trade-offs, where the brain prioritizes neural integrity over peripheral lipid balance. These lipidome changes and redox-driven shifts threaten both individual health and ecosystem stability, emphasizing the need for policies to regulate pesticide use and minimize exposure risks.

Graphical abstract

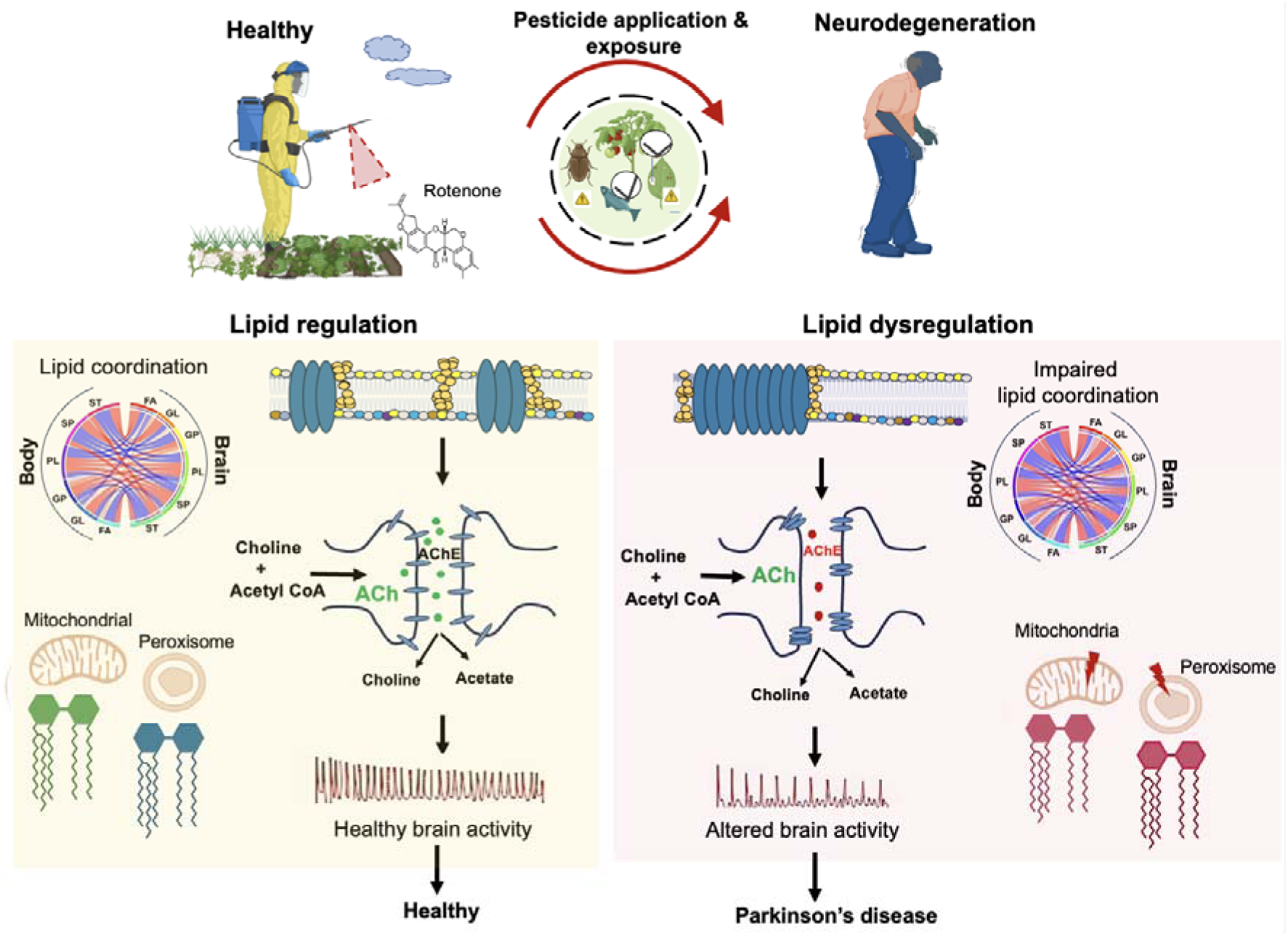

## 1. Introduction

With the global rise in aging populations, the increasing incidence of neurodegenerative diseases highlights a growing concern. Pesticides are increasingly recognized as environmental contributors to brain disorders, impacting cellular function and brain health. Among these, rotenone stands out as a key environmental stressor associated with the development of neurodegenerative diseases. Rotenone, a biopesticide, is a naturally occurring lipophilic compound obtained from roots and stems of certain tropical and subtropical plants like species of *Lonchocarpus* and *Derris* is globally used in agriculture to enhance crop yield [1,2]. The use of rotenone as a pesticide expanded globally, especially with the rise of organic farming systems and agricultural practices, due to its natural insect control capabilities [1,3]. Besides that, it is also used to control the over-invasions of species in the freshwater ecosystems and to manage fish population [3]. The major mechanism of action for this pesticide has been reported to occur through interference with cellular respiration by inhibiting mitochondrial complex I, NADH-ubiquinone oxidoreductase complex, subsequent depletion of ATP, energy in the cell and thereby causing the death of target insect [4].

Being a plant-derived natural pesticide, rotenone was initially considered a safer alternative to synthetic pesticides, for its wider application in organic farming systems. Later, it was reported that rotenone exposure has broader environmental impact and potential toxicity to humans and non-target organisms, became a growing concern due to its non-selective action [5,6]. Due to the potential non-target effects, led to increased regulation in many countries including United states and European Union. The World Health Organization (WHO) categorizes it as a “Class II hazardous” pesticide. The United States of environmental protection agency (USEPA) allows the application of rotenone in water bodies at specific concentrations, with stricter limits in recreational areas and sewage systems due to its toxicity [7]. Even at low concentrations, significantly effects wide range of organisms including beneficial insects, aquatic species and terrestrial wildlife [8, 9]. It can be absorbed through dermal contact, inhaled as aerosol or dust or ingested via water or dietary contamination by rotenone [5]. The long-term health impacts of chronic rotenone exposure have sparked the most concerns in the recent years [9,10].

Humans are commonly exposed to rotenone, which degrades slowly in fruits and vegetables, often exceeding the permitted threshold of 0.04 mg·kg ¹. In processed vegetables, the rotenone levels can be up to 4.8 times higher than in raw vegetables. This underscores the risks of rotenone exposure through food and water. Epidemiological studies identified the association between rotenone exposure and Parkinson’s disease (PD) [10,11]. Agricultural workers, pesticide applicators, individuals living near areas where rotenone is heavily used and through environmental exposure have been shown to have a higher incidence of PD [10,11,12,9]. A case-control study by U.S. Agricultural Health study (AHS) reported a 2.5-fold increased risk of PD among individuals who used rotenone, with longer durations of use further amplifying the disease risk [10]. A more recent and comprehensive nested study within the French agricultural cohort, AGRICAN, revealed that exposure to pesticides including rotenone increased the risk of PD by up to 1.6-fold, and 1.2% of participants in this cohort were reported to have PD [11]. Although the exact cause of PD remains unidentified, rotenone has been strongly implicated as causative factor in previous cohort studies.

The growing body of evidence linking rotenone to PD has significant implications for health, occupational safety and environmental policy [5,10,11,12,9]. The USEPA had recently identified risks of concern with rotenone application and recommend the need for the environmental health risk assessment of this pesticide to evaluate the potential risks pose to human health and environment [5]. Alternative systems to humans are indispensable in risk assessment of pesticides, as in the present case rotenone induced health risk associated pathogenesis, PD [2]. These animal models enable researches to explore the biomarkers and cellular process involved in chronic rotenone induced PD by providing valuable insights that would be difficult to achieve through human studies alone [2,13]. Recent rotenone induced animal model study identified that it reflects accurate human clinical situation by delayed accumulation of alpha-synuclein in neurons of the substantia nigra [14]. Furthermore, a recent study identified that exposure to rotenone in developing zebrafish embryos induces features commonly associated with PD pathology [15].

The brain tissue plays a major role in rotenone associated PD development as its exposure impairs the ability of mitochondria to buffer calcium, and, excessive intracellular calcium exacerbates oxidative stress and promote mitochondrial membrane permeability transition [15,16]. Additionally, rotenone induced mitochondrial dysfunction can disrupt the metabolism leading to imbalances in lipid pathways that regulates membrane fluidity, cellular homeostasis, cell survival and apoptosis [17,18]. Although lipid levels are altered in various conditions of metabolic stress, very little is known about their role in brain and systemic energy metabolism in response to rotenone exposure. Furthermore, the lipid biomarkers associated with in vivo chronic exposure to rotenone that lead to PD are largely unknown. Therefore, the system-wide lipid profiling of rotenone-exposed animals would not only allow us to understand the biomarkers for environmental risk assessment but also a larger landscape of lipid homeostasis in PD progression.

Here we hypothesized that chronic environmental concentration of rotenone exposure alters the metabolic landscape leading to lipidome dysregulation in the brain and progressive PD-like phenotype. We use *Drosophila melanogaster* to investigate the health risk assessment of rotenone by characterizing lipidome alteration in brain and peripheral tissues. *Drosophila* has a remarkably well-conserved biological system that shares many fundamental features of humans. About 75% of human disease-causing genes have functional counterparts in *Drosophila*, including those involved in neurological disorders. Using untargeted liquid chromatography-high-resolution accurate mass spectrometry (LC-HRAMS)-based lipidomics, we characterized lipidome alterations in brain and peripheral tissue of *Drosophila melanogaster* under basal and environmentally relevant concentration exposure to rotenone. In particular, 414 lipids structurally identified with significant dysregulation of glycerophospholipids (GPs), sphingolipids (SPs), fatty acyls (FAs), and Glycerolipids (GLs) under rotenone exposure. Furthermore, the biomarker lipids associated mitochondrial and peroxisome dysfunction related to PD caused by rotenone were identified. More importantly, the redox imbalance in lipid profile was observed. Moreover, behaviour alterations, biochemical perturbations were investigated.

## 2. Material and methods

### 2.1 Chemicals and reagents

Rotenone (> 99% purity; Tokyo chemical industry, Japan), Acetonitrile (Optima LC-MS grade, Sigma Aldrich), methanol, ethanol, isopropanol (LC-MS grade, Sigma Aldrich, St. Louis, MO, USA), Ultra-pure water (LC-MS grade, Sigma Aldrich, St. Louis, MO, USA). Sodium phosphate (NaHPO3) (Sigma Aldrich, St. Louis, MO, USA), 4-(2-Hydroxyethyl) piperazine-1-ethanesulfonic acid (HEPES), Sodium lauryl sulfate (SDS) and Amplex@Red Acetylcholinesterase Assay Kit was procured from ThermoFisher Scientific, USA. Hydrogen peroxide (H2O2; Himedia), Dichloro-dihydro-fluorescein diacetate (DCFH-DA; Cayman chemicals), Thiobarbituric acid (TBA), Tetraethoxy propane (TEP) (Tokyo chemical industry, Japan). Superoxide Dismutase Activity Kit (Abcam Japan).

### 2.2 Fly lines and maintenance

The wild-type *Canton-S* strain of *D. melanogaster* was maintained on a 4% sugar cornmeal-yeast-agar medium at 24 ± 1 °C in humidity (60-70%) with 12-hour light/12-hour dark cycle at 400 lux. Male flies were segregated and kept on the control diet for 4-5 days. Five-days-old male flies were used to proceed with the experiment.

### 2.3 Rotenone exposure using *Drosophila melanogaster*

For the determination of appropriate concentrations and exposure period for *D. melanogaster* treatment, the rotenone compound was dissolved in 0.1% dimethyl sulfoxide (DMSO) according to previous the protocol described by Nazir et al. Based on survival and longevity data, a 7-day exposure period was selected to assess the individual effects of rotenone. The experimental group was a control group (flies fed agar-sucrose media containing 0.1% DMSO) and three groups of rotenone treatment included 125 µM, 250 µM, and 500 µM. The rotenone was uniformly mixed with 5 mL of agar-sucrose media for each group. Rotenone concentration of **500 µM in 5 ml** of food corresponds environmental exposure levels and allows for the modeling of rotenone’s toxicological effects in controlled conditions [43]. Male flies were collected immediately after eclosion and kept under a 12-hour light/12-hour dark cycle at 25°C. Five-day-old flies were considered for the treatment. Before being transferred to the rotenone-containing media, flies were kept for 5-hour starvation in empty vials. After 7 days of exposure, the flies were euthanized, and their brain and body tissues were quickly processed separately to analyze global lipidome and oxidative stress markers.

### 2.4 Determination of locomotor activity using *Drosophila* activity monitor

To assess the effects of rotenone treatment throughout light-dark (LD) cycles, locomotor activity was measured with Drosophila Activity Monitors (DAMs, TriKinetics). Adult male flies were momentarily anaesthetised with CO2 and placed in 65 mm-long glass tubes (TriKinetics) with one end having a feeding supply (5% sucrose + 1.2% agar) and the other containing a cotton plug. The locomotor activity was continuously recorded over 10 days using the DAM2 system (TriKinetics) to assess free-running locomotor rhythms. Data were collected in 30-minute intervals and analyzed using the GraphPad Prism. Throughout the experiment, flies were maintained under white LED lighting at approximately 400 lux in a temperature-controlled incubator set to a 12-hour LD cycle at 25°C. Each experiment included a minimum of 8 flies per condition.

### 2.5 Negative geotaxis assay

The negative geotaxis experiment, which evaluates fruit flies’ innate escape behaviour, is a frequently utilised approach in research on themes ranging from metabolic processes to neurodegenerative illnesses (19). This test was used in this investigation to evaluate the locomotor activity of control and rotenone-treated flies in accordance with known protocols [20]. A 25 mL graduated glass cylinder (18 × 2 cm) was used to hold groups of 20 flies. After a 1-minute acclimatisation period, the flies were tapped to the cylinder’s bottom, and the number of flies that ascended beyond or stayed below the 10-centimeter mark was measured after 30 seconds. This treatment was done five times with a one-minute rest period between each. The climbing index for each experiment was computed using the following formula

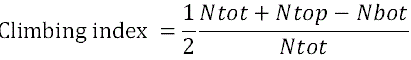

Ntop = number of flies climbing above 10 cm

Nbot = those remaining below 10 cm

Ntot = total flies number

The data were reported as mean ± SEM from six independent biological replicates.

### 2.6 Survival assay

The survival of experimental flies was assessed as previously described, with minor alterations. [21]. The effect of rotenone on the survival of *D. melanogaster* was evaluated over a seven-day treatment period. Each experimental group i.e., the control and rotenone treated was monitored daily by by measuring the number of viable flies. Fifty adult flies were randomly assigned to each replicate at the start of the treatment. To avoid confounding effects from handling, no food was replaced during the treatment period. The experiment was conducted in triplicate to ensure reproducibility and reliability of the results.

### 2.7 Determination of dopamine level

The extracted brain tissue of *D. melanogaster* was resuspended with 200 μl of 80% chilled methanol. LC-MS-based selective ion monitoring methods (SIM) analysis of dopamine was determined by using a Vanquish UHPLC-Liquid Chromatography (LC) system coupled to a Orbitrap 240 mass spectrometer (Thermo Scientific). LC separation was performed by using ACQUITY UPLC BEH Amide (2.1x 150 mm 1.7-micron particle size; Acquity Waters) column with a gradient solvent A (0.1% formic acid in water); and solvent B (acetonitrile with 0.1% formic acid). A 22.200 min elution gradient of solvent A:0.1% formic acid in water and Solvent B: acetonitrile was used. The gradient as follows: 0 to 2 min 80% B; 17 min 20%B; 17.100 min to 22.100 min 80% B followed by 80% B till 22.200 min. MS parameters were: spray voltage 3.5□kV for positive and probe temperature 330 °C; sheath and auxiliary gases were 35 and 10 arbitrary units, respectively; and mass of dopamine 154.0861 m/z was set for SIM data acquisition and settings of AGC target and resolution as Balanced 120000 respectively.

### 2.8 Tyrosine hydroxylase (TH) activity

TH activity was measured using the procedures described previously by Vermeer et al. [22] and Musachio et al. [23]. Flies were cryoanesthetized and decapitated, with 20 brains per group collected for each experiment. The collected brains were homogenised in 250 μL of Tris-HCl buffer (0.05 M, pH 7.2), then centrifuged at 13,000 g for 5 minutes at 4°C. A 100 μL sample of the supernatant was combined with an equal volume of a reaction solution M tyrosine, and 200 μM sodium periodate. The enzymatic process was monitored spectrophotometrically at 475 nm. The results were normalised and expressed as a percentage reduction from the control. To ensure accuracy and reproducibility, five independent biological replicates (n = 5) were performed.

### 2.9 Extraction of rotenone and its metabolite in tissues for LC-MS

Dissected tissue was homogenized with one milliliter of 100% chilled Ethanol using a *Drosophila* tissue homogenizer (pellet pestle mortar, Kontes) using the previously described extraction methods with modification [24]. Tissue was sonicated for 5 minutes and 25 minutes vortex using a multitube vortexer (iGENE LABSERV) followed by 15 minutes of centrifugation by maintaining a 4^0^ C temperature at 14000 rpm. The extraction process was repeated three times similarly.

### 2.10 LC-MS-based determination of rotenone and its metabolites in brain tissues

The extracted tissue was reconstituted with 200 l of 100% cold ethanol. LC-MS analysis was performed using a Vanquish UHPLC-Liquid Chromatography system paired with an Orbitrap 240 mass spectrometer. Rotenone and its metabolite were identified using an optimized LC-MS method with the following conditions: a C18 column (2.1 x 150 mm, 1.8-micron particle size; Zorbax Eclipse Plus C18) was employed for LC separation. The mobile phase consisted of solvent A (0.1% formic acid in water) and solvent B (acetonitrile with 0.1% formic acid). The gradient profile was as follows: from 0 to 3 min, 5% B; at 13 min, 50% B; at 20 min, 75% B; at 26 min, 75% B; at 28 min, 5% B; and from 28 to 30 min, 5% B. l/min, an injection volume of 3 μl, a column temperature of 40°C, and an autosampler temperature of 4 °C. MS (Product ion scan) was performed in positive mode with normalised collision energies of HCD at 30 and 60.

### 2.11 Lipid extraction and sample preparation for LC-MS-based lipidomics

The brain and body tissues of dissected flies were homogenised with 100□µl of cooled 1XPBS using a handheld motorised pestle. Lipids were extracted using a modified method from Xue Li Guan et al. (25). In summary, 500□µl of chloroform:methanol (1:2, v/v) was added to the homogenate, vortexed for one minute, and then placed on an Eppendorf thermomixer (Thermomixture C), stirring at 1,000 rpm for 2 hours at 4°C. After adding 200 μl of chloroform and 200 μl of water to the mixture, the phases were separated by centrifugation (Eppendorf, Centrifuge 5424R) at 9,000 rpm for 2 minutes at 4 °C. The lower organic phase was recovered and dried using a vacuum concentrator before lipidome analysis. The lipid extracts were spiked with internal standards and LC-MS analysis was performed as previously described [26].

### 2.12 Untargeted lipid profiling of tissues using high-resolution accurate LC-MS

The extracted brain and body tissue lipid samples were reconstituted with 200 μl of chilled absolute Acetonitrile: methanol (1:1 v/v) and lipids are separated by using the reverse phase chromatography with sight modification as described previously [26]. Lipidome analysis was conducted by using a Vanquish UHPLC system coupled to an Orbitrap 240 mass spectrometer (Thermo Scientific). The LC separation was performed by using the previously described methods [27]. Briefly, C18 column (2.1x 150□mm 1.8-micron particle size; Zorbax eclipse plus C18) with a gradient of solvent A (Acetonitrile: Water, 60:40 v/v) and solvent B (Isopropanol: Acetonitrile, 90:10 v/v). Gradient solvents were consisting of solvent A (Water: Acetonitrile, 40:60 v/v) and solvent B (Acetonitrile: Isopropanol, 10:90 v/v) with 10mM ammonium acetate and 0.1% formic acid. A 27.00 min elution gradient of solvent A and Solvent B was used. The gradient is as follows: 0 to 3 min 30% B; 5 min 43%B; 5.100 min 55%B; 10 min 65 %B; 12.500-13.500 min 85% B; 18 min to 23 min 100%B followed by 30% B till 27.00 min. Other LC parameters were a flow rate of 300 μl/min, a column temperature of 40°C, an injection volume of 2 μl, and an autosampler temperature of 4°C. MS was performed with a HESI II probe in both positive and negative polarity. The MS parameters are as follows: Full scan range: 100 to 1500 m/z with AGC target and resolution set to Balanced 120000, respectively; spray voltage: 3.8 kV and 3.5 kV for positive and negative modes; probe temperature: 330 °C; sheath and auxiliary gases: 35 and 10 arbitrary units, respectively. Xcalibur 4.5 (Thermo Scientific) software was used to record the data. Additionally, lock-mass correction was conducted using common low-mass contaminants in every analytical run to improve calibration stability. HCD collision energies 20, 35, and 60 were used as the normalized collision energies for data-dependent analysis. Lipids were identified using LIPIDMAP [28], lipid home, LipidBlast and *Drosophila* lipidome source file**s** [29].

### 2.13 Preparation of homogenates for biochemical assays

After seven days of rotenone treatment, flies were anesthetized and euthanized on ice. The flies were then dissected into brain and body tissues. For each oxidative stress assay, the required number of brains and bodies were pooled and homogenized following the specific protocol for each analysis. The biochemical assay activities were measured separately in the brain and body tissues. Calculating the enzyme activity involved normalization to the total protein content. Total protein content was estimated in samples using Bradford method having bovine serum albumin as the standard.

### 2.14 In vivo ROS imaging

ROS detection was carried out using 2,7’-dichlorofluorescein (H2DCF) according to the methodology reported by Vaccaro et al., 2024 [30]. Flies were anaesthetised with cold and dissected into brains in 1X phosphate-buffered saline. The tissues were treated on a shaker at room temperature with 10 µM H2DCF for 10 minutes in darkness. Following incubation, the tissues were rinsed three times in PBS for 5 minutes each, while shaking at room temperature. The samples from each independent replication were then put on a glass slide with glycerol and photographed immediately using consistent settings on a Leica SP8 confocal microscope with a 488 nm excitation wavelength. The obtained photos were analysed with ZEN software.

### 2.15 ROS quantification

The oxidation of 2,7-dichlorofluorescein diacetate (DCFDA) was measured in the sample supernatant as an indication of free radical formation and oxidative stress, using the procedure reported by Pérez-Severiano et al. [31]. Twenty fly brains were homogenised in 100 μL of buffer (pH 7.4) and centrifuged at 8,000 × g for 5 min. A 10 μL sample of the L of 1X phosphate-buffered saline (pH 7.0) and 1 μL of 5 mM DCFDA. After 30 minutes of incubation in the dark, the fluorescence emission of DCFH produced by the oxidation of DCFDA was measured using a spectrophotometer with an excitation wavelength of 492 nm and an emission wavelength of 517 nm. The results were presented as fluorescence intensity per mg of protein. Data were averaged from five independent biological replicates, with each group consisting of 100 flies.

### 2.16 Estimation of lipid peroxidation (LPO)

Tetraethoxypropane was employed as an external reference for determining malondialdehyde (MDA) levels during LPO [32]. To prepare the mixture, 25 μL of 10% brain homogenate was mixed with 150 μL of 10% SDS, 1.0 mL of 0.8% thiobarbituric acid, and 1.0 mL of 20% acetic acid solution (pH 3.5). The resulting mixture was then incubated in boiling water at 100°C for one hour. After incubation, the coloured substance was extracted with 3.0 mL of normal butanol, and the absorbance was measured at 535 nm. Normal butanol was used as a blank. The analysis was carried out in five independent replicates, with each sample analysed twice. The MDA concentration was measured and expressed as nanomoles per milligram of protein.

### 2.17 Catalase activity assessment

Catalase activity was measured according to the methodology published by Anet et al. [33]. This assay measures the enzyme’s ability to break down H□O□. To begin, make solutions of dichromate acetic acid, 0.02 M H□O□, and 0.01 M sodium phosphate buffer. The assay procedure involved adding 1 mL of the buffer to a 15 mL Falcon tube, followed by the addition of deionized water (dH□O). Next, 25 μL of a 10% homogenate was introduced into the tube. After mixing, 0.5 mL of 0.02 M H□O□ was added, and the mixture was vortexed again. Then, 2.0 mL of dichromate acetic acid was added, and the mixture was incubated in boiling water at 100°C for 15 minutes. The absorbance of the resulting colored product was measured at 570 nm using a spectrophotometer. The assay was carried out in five independent experiments, each with triplicate samples. Results were normalised for protein concentration and represented as micromoles of H□O□ decomposed per milligramme of protein (µmol H□O□ /mg protein).

### 2.18 Superoxide dismutase activity (SOD)

SOD activity in Drosophila tissues exposed to rotenone was measured using a biochemical kit from Abcam [34]. Brain and body tissues were homogenized in ice-cold 0.1 M Tris buffer (pH 7.4) containing 0.5% Triton X-100, 5 mM β-mercaptoethanol, and 0.1 mg/mL phenylmethylsulfonyl fluoride using a pestle and mortar. The homogenate was centrifuged at 14,000 × g for 5 minutes at 4°C, and the supernatant was collected while maintaining it on ice. The assay was performed by adding 200 μL of the working solution to each sample well, and 20 μL of dilution buffer was added to the blank wells. Then, 20 μL of the sample was added to the assay wells, followed by 20 μL of enzyme working solution. The reaction mixture was incubated at 37°C for 20 minutes. Optical density was measured at 450 nm. SOD activity was calculated as the percentage of inhibition using the formula:

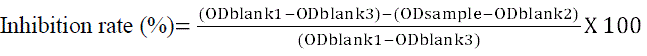

### 2.19 Determination of acetylcholinesterase (AChE) activity

Twenty brains and twenty bodies from each group were homogenized separately with 100 μL of homogenizing buffer (pH 7.4), followed by centrifugation at 8,000×g for 5 minutes at 4°C, according to the protocol described by Musachio et al. [35]. For the analysis, 15 μL of the supernatant from both the brain and body samples were diluted with 1X reaction buffer. Two positive controls were prepared: one with 0.2 U/mL AChE and the other with 20 mM H□O□. All necessary reagents were added to prepare the working solution for the sample reaction. The samples were incubated with the working solution for 30 minutes or longer at room temperature in the dark [36]. The AChE activity was measured spectrophotometrically [36].

### 2.21 Statistical analysis

Rotenone and its metabolites were identified using Qualbrowser (Xcalibur 4.5 software). For lipid analysis, the raw data were processed using MSDIAL version 4.9.2. Lipid identification was performed using the LipidBlast database. All steps in the MSDIAL lipidome data processing were carried out with previously described parameters, with minor modifications [37]. Multivariate statistical analysis was performed using Principal Component Analysis (PCA) and Partial Least Squares Discriminant Analysis (PLS-DA) to assess group differences. The PLS-DA model was built with R software packages. A heatmap was generated based on the abundance of differential lipids using R software. Correlation analysis was used to calculate the correlation coefficient (r) between different lipid groups. The following criteria were applied for univariate analysis to identify differential lipids: the Variable Importance for the Projection (VIP) in the first two Principal Components (PCs) of the PLS-DA model was set to ≥ 1. Fold Changes (FCs) were categorized as upregulated if FC ≥ 1 and downregulated if FC ≤ 1. A P-value of less than 0.05 was considered statistically significant. Differentially expressed lipids were defined by VIP ≥ 1, P-value < 0.05, and FC ≥ 1 or ≤ 1. Statistical analysis and data visualization were performed using GraphPad Prism version 8.1.

## 3 Results

### 3.1 Chronic rotenone exposure induced behaviour abnormalities and dopaminergic neurodegeneration

We initially assessed the impact of prolonged rotenone exposure on the behavior *of Drosophila melanogaster* using a commonly employed test that evaluates phenotype consistency with neurodevelopmental impairments (Fig. 1A). Three distinct concentrations of rotenone (125 µM, 250 µM, and 500 µM) were administered through dietary exposure to represent varying chronic doses. In the negative geotaxis test, a significant number of rotenone-exposed flies failed to climb 15 cm within 30 seconds compared to the control group: 22% for the 125 µM, 36% for the 250 µM, and 44% for the 500 µM groups (Fig. 1B); statistical analysis showed significant differences for all concentrations (*P* < 0.0003). These results suggest a reduction in overall activity and a notable decline in locomotor behavior in the rotenone exposed flies. As neurodevelopmental disruptions are often linked to sleep disturbances, we further investigated whether rotenone exposure affected circadian rhythms. We continuously monitored the activity-rest cycles of fruit flies under standard light-dark (LD) conditions over seven days with varying concentrations of rotenone. A dose-dependent reduction in total daily activity was observed in rotenone-treated flies compared to controls (Fig. 1C): 125 µM (3-fold decrease, P = 0.002), 250 µM (3.5-fold decrease, *P* = 0.0014), and 500 µM (4-fold decrease, *P* = 0.0010). Additionally, rotenone-exposed flies showed a shift in their activity patterns, with a delayed reduction in behavior prior to the onset of the dark period. Specifically, these flies were less active on average one hour before the dark period (P < 0.05). To further investigate the impact of rotenone on daily activity patterns, we examined the 24-hour activity cycle and noted significant differences between control and rotenone-exposed groups, with a marked reduction in activity during the light phase compared to the dark (P < 0.05). Furthermore, a phase shift in the circadian rhythm was observed at three hours before the dark period (CT9). These findings collectively demonstrate that chronic exposure to rotenone adversely affects both locomotor and circadian rhythms (Fig. 1D). The actogram analysis on the seventh day revealed a decrease in activity for flies exposed to rotenone at all concentrations when compared to controls (Supplementary Fig. S1). Chronic rotenone exposure significantly affects the survival of *Drosophila* (Fig. 1E). The declined locomotor and circadian activity behavior in rotenone-treated flies directly reflects the loss of dopaminergic neurons in the brain. As these neurons die, the tyrosine hydroxylase expression diminishes. To confirm the same, tyrosine hydroxylase was measured in rotenone-exposed *Drosophila* brains. Tyrosine hydroxylase levels were significantly lower at 125 µM (*P* = 0.0074), 250 µM (*P* = 0.0133), and 500 µM (*P* = 0.0110) of rotenone as compared to controls (Fig. 1F). Further, in neuronal degeneration conditions, the inhibition of tyrosine hydroxylase activity directly impairs dopamine production whereas in normal conditions dopamine exerts feedback inhibition on tyrosine hydroxylase through its metabolites. To identify the impact of rotenone on the dopaminergic system, dopamine (DA) levels were measured using LC-MS. DA levels were significantly diminished in the rotenone-exposed groups compared to the controls 125 µM (*P* = 0.0033); 250 µM (*P*< 0.0001); 500 µM (*P* < 0.0001) (**Fig. 1G**). This reduction in DA levels was consistent with the decrease in TH levels and directly linked to impaired dopamine feedback regulation. These findings suggest that rotenone exposure induces impairment of the dopaminergic system in this model.

**Figure 1:**
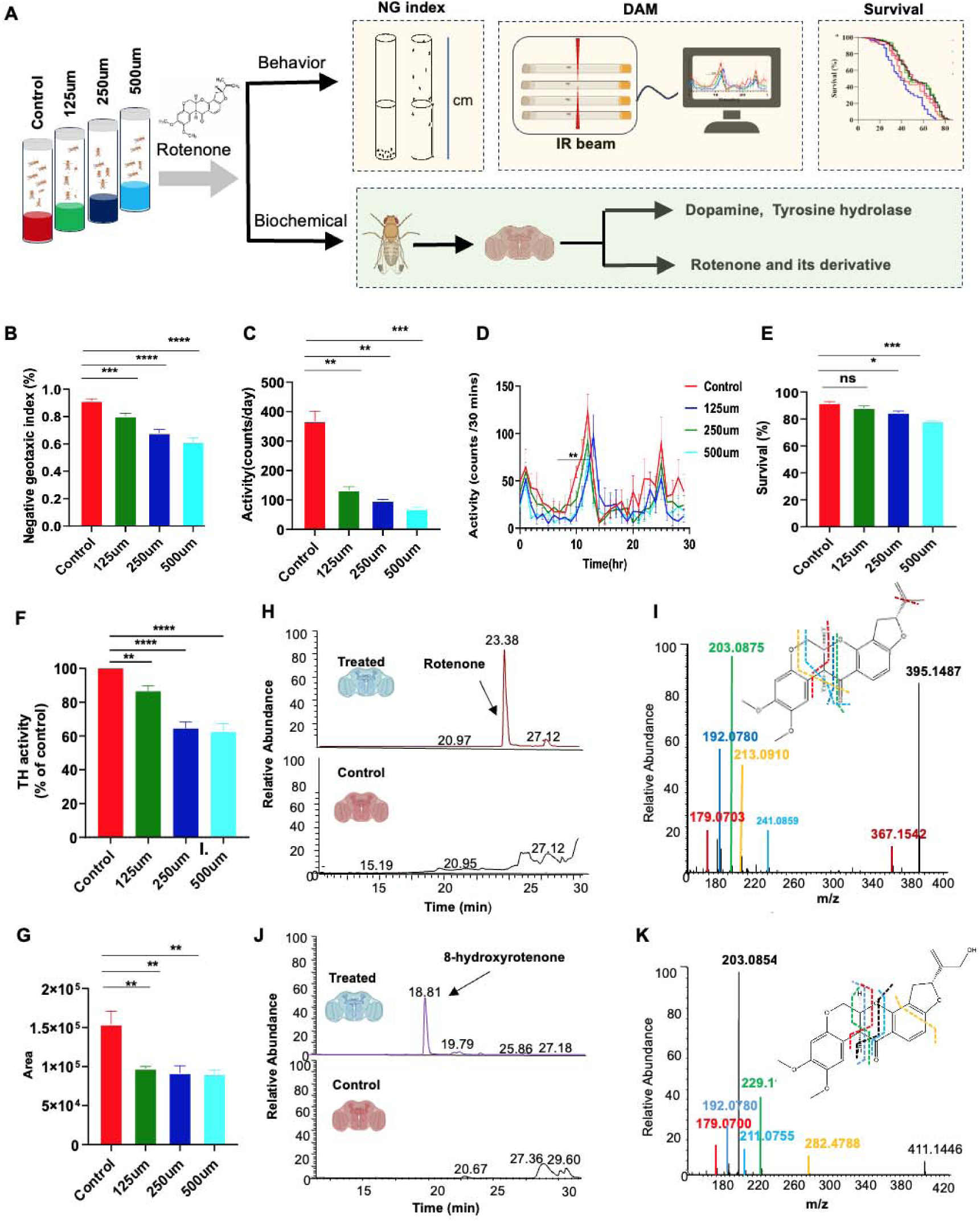
Rotenone-induced significant alteration of behavior and tissue specific metabolites. **A.** Schematic representation of experimental plan for behavior analysis **B.** Negative geotaxis index reduced in rotenone-exposed flies. N = 6 with twenty flies per group **C.** Total Activity per day **D**. Activity pattern reduced during Rotenone treatment. **E.** Survival rate of flies after the seven days of rotenone treatment. **F**. Tyrosine hydrolase activity in *Drosophila* brain tissue **G.** Dopamine levels in brain tissues of control and rotenone treated flies. **H, I.** TIC and MS2 spectra of rotenone in the brain tissue **J, K.** TIC and MS2 spectra of rotenone metabolites (8-hydroxy rotenone) in the brain tissue. Values are expressed as Mean□±□ Standard Error of Mean (*n*□=□5). Significant differences from the control are indicated p* < 0.05, p**<0.005, p****<0.0001 Student’s t-test.

### 3.2 LC-MS/MS analysis identifies rotenone and its metabolites in brain tissue under chronic exposure of rotenone

To ascertain the intake of rotenone by exposure, LC-MS-based assay was used to detect brain rotenone following chronic exposure. Rotenone and its metabolites were analyzed in *D. melanogaster* brain tissues [24] following exposure to rotenone. We have detected rotenone (Figs. H & I) and its metabolite 8-hydroxy rotenone in the brains of *D*. melanogaster (Figs. J & K). Significant accumulation of rotenone (m/z =395.1487), 0.91ng/brain was observed in brain tissue. Furthermore, the identification of its primary metabolic product, 8-hydroxy rotenone (m/z =411.1446).

### 3.3 Rotenone exposure induces lipidome alterations

#### 3.3.1 Chronic rotenone exposure alters the abundance of lipid classes

Next, we asked how *Drosophila* lipidome varied in different tissues including brain and body, after chronic exposure to rotenone at different concentrations. To answer this, we have applied lipidomics profiling using high-resolution accurate mass spectrometer Orbitrap exploris. Total extracts were analyzed using this LC-HRAMS in DDA mode (Fig. 2A). Tight clustering of pooled QC samples explains the good reproducibility (Fig. 2B; Fig. S2). Lipids were identified by matching their masses at sub-p.p.pm. accuracy having elemental composition constraints of a specific class [38,39]. Furthermore, peak assignments were validated by MS/MS fragmentation performed on the respective precursors. Of total, we systematically identified 414 lipids (Figure 2C) from six lipid categories including — Fatty Acyls (FAs), Glycerolipids (GL), Glycerophospholipids (GP), Prenol Lipids (PL), Sphingolipids (SP), and Sterol Lipids (ST) (Figure 2D) in both tissues. Scatter plots show the range of lipids contained in each tissue in both ESI (+) and ESI (-) ionization modes. A large group of glycerophospholipids were found in brain tissue in ESI (-) ionization mode (Fig. S3A), whereas high levels of glycerolipids and fatty acyls were found in ESI (+) ionization mode (Fig. S3B). In case of *Drosophila* body tissue, relatively large group of glycerophospholipids and glycerolipids were found in ESI (-) (Fig. S3C) and ESI (+) ionizations (Fig. S3D) respectively. To understand how these six lipid categories altered in response to rotenone in both brain and body tissues, pie charts were constructed (Fig. 2E). The total percentage of GP lipids are found to be significantly reduced (16% and 7% for 125uM, 16% and 10% for 250uM, and 14% & 8% for 500uM rotenone exposure) in both tissues with varying concentrations of rotenone exposure as compared to control lipid levels in the brain (17%) and body (12%) tissues. Similarly, the abundance of other classes of lipids including GL, FA, SP, PL, and ST are significantly altered in response to different concentrations of rotenone exposure. Additionally, of total, we identified 27 lipid subclasses in brain and 29 lipid subclasses abundance in body tissue. The lipid subclasses including acylcarnitine (CAR), free fatty acids (FFA), lysodiacylglyceryltrimethylhomoserine (LDGTS), N-acyl ethanolamines (NAE), N-acyl glycyl serine (NAGly), N-acyl taurine (NATau), N-acyl ornithine (NAOrn), Diacylglycerol (DG), ether-linked diacylglycerol (Ether-DG), triacylglycerol (TG), ether-linked triacylglycerol (Ether-TG), cardiolipin (CL), lysophosphatidylcholine (LPC), lysophosphatidylethanolamine (LPE), Lysophosphatidylglycerol (LPG), lysophosphatidylinositol (LPI), phosphatidylcholine (PC), Ether-linked phosphatidylcholine (Ether-PC), phosphatidylethanolamine (PE), ether-linked phosphatidylethanolamine (Ether-PE), phosphatidylglycerol (PG), phosphatidylinositol (PI), phosphatidylserine (PS), coenzyme Q (CoQ), ceramides (CER), hexosylceramides (HexCer), Taurocholic acid (TCA), sterol derived lipids (ST), ether-linked phosphatidylinositol (Ether-PI), ether-linkedlysophosphatidylethanolamine (Ether-LPE) (Fig S4).

**Figure 2:**
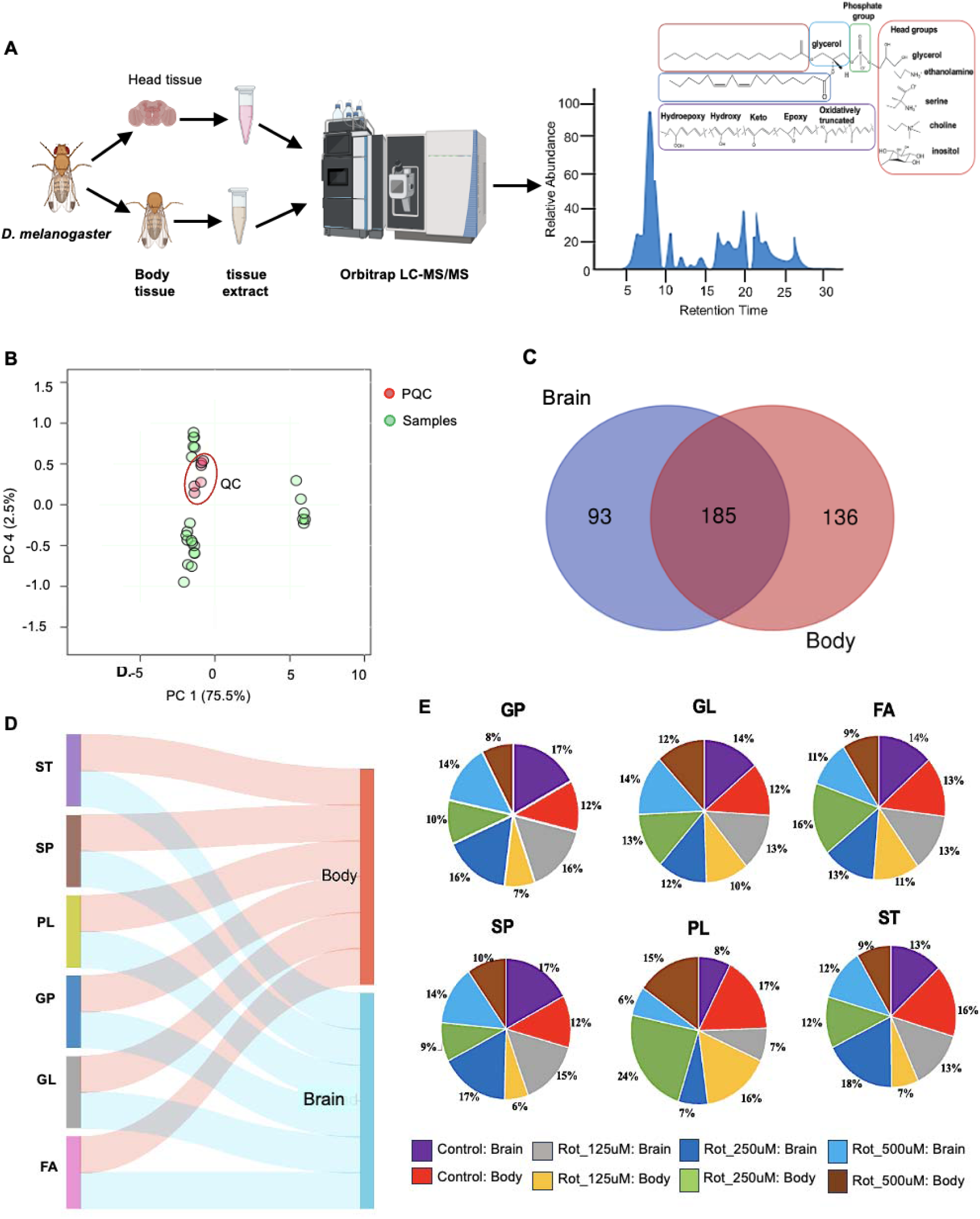
Rotenone induced lipidome alteration in brain and body tissue. **A.** Lipidomics study design summary **B.** PCA plot for pooled QC and Samples **C.** Venny diagram representation of total lipids in both tissues **D.** Sanky dot plot showing lipid category in brain and body tissue **E.** 2D pie chart showing the percentage of lipids in brain and body tissue with varying concentration of rotenone. Rot: Rotenone concentrations 125, 250 and 500uM

#### 3.3.2 Chronic rotenone exposure alters *Drosophila* brain lipidome

To simplify the dataset and map it into a low-dimensional space, multivariate analysis was employed to distinguish the metabolic profiles across sample classes. Initially, unsupervised principal component analysis (PCA) was applied to data obtained from both ESI (-) and ESI (+) modes. The resulting PCA score plots illustrated the distribution of samples, where clustered points represented similar metabolomic profiles, while dispersed points indicated distinct metabolomic differences. The PCA analysis of the mass spectral data for all samples provided insights into the overall structure of the dataset. In this analysis, the first two PCs represented for 85.9% of the variance in ESI (-) mode and 77.1% in ESI (+) mode (Fig. 3A & B). Specifically, in ESI (-) mode, PC1 captured 75.6% of the variance, while PC2 accounted for 10.3%. Similarly, in ESI (+) mode, PC1 explained 64.8% and PC2 captured 12.3% of the total variance. The PCA scores plot revealed four distinct clusters. A notable separation (P < 0.05) was observed between control and rotenone-treated *Drosophila* along the first principal component. Figures S5A and B shows individual and cumulative explained variances for primary components.

**Figure 3:**
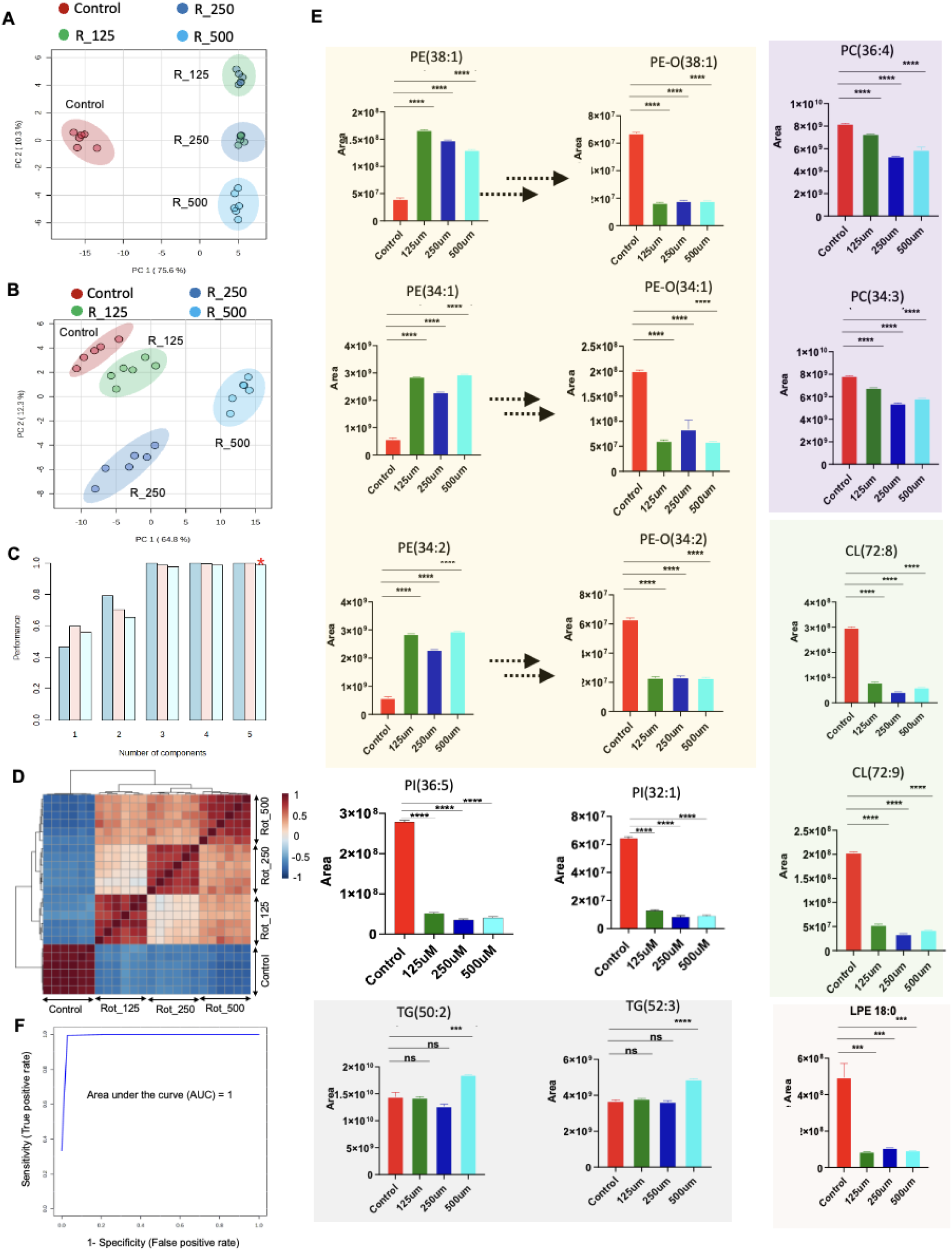
Significantly altered marker lipids in brain tissue under chronic exposure of rotenone. A & B. Two principal components accounted for 85.9% of the variance in ESI (-) mode and 77.1% in ESI (+) mode C. For ESI (-ve), PLS-DA demonstrated strong performance with cumulative values of R² = 0.99712, Q² = 0.98678, and Accuracy = 1 D. Pearson’s correlation E. Marker lipids including PE(38:1), PE(34:1), and PE(34:2); PE O-(38:1), PE O-(34:1), and PE O-(34:2); PI(36:5) and PI(32:1); TG(50:2) and TG(52:3); PC(36:4) and PC(34:3); CL(72:8) and CL(72:9); and LPE 18:0 F. ROC curves distinguished rotenone-exposed *Drosophila* from controls.

PLS-DA was conducted to pinpoint the metabolites responsible for the observed separation between control and rotenone-exposed Drosophila in both ESI (-) and ESI (+) modes (Figs. S5C & D). The scores plots revealed distinct clustering of individual samples within their respective groups for both ionization modes. However, visual inspection of PLS-DA scores plots alone is insufficient to determine predictive power. To address this, an internal cross-validation was carried out. For ESI (-) mode, the PLS-DA demonstrated strong performance with cumulative values of R² = 0.99712, Q² = 0.98678, and Accuracy = 1 (Fig. 3C). Similarly, the ESI (+) mode model showed R² = 0.99742, Q² = 0.98537, and Accuracy = 1 (Fig. S6A). Permutation testing was then conducted to assess the statistical significance using 500 permutations. The analysis confirmed the high predictive power of the optimal models for distinguishing rotenone-induced chemical toxicities from controls, with significance determined at P < 0.01 (Figs. S6B & C). To compare control and rotenone-treated brain sample extracts, Pearson correlation was used, showing a strong positive correlation between each sample group and its corresponding control and treatment group (Fig. 3D). Based on VIP values (VIP > 1) from the PLS-DA model, 15 lipids were identified as key differentiators (Table 1). These included phosphatidylethanolamines (PE) such as PE(38:1), PE(34:1), and PE(34:2); plasmalogen or ether-linked PE (PE O-) lipids such as PE O-(38:1), PE O-(34:1), and PE O-(34:2); phosphatidylinositols (PI) including PI(36:5) and PI(32:1); triglycerides (TG) such as TG(50:2) and TG(52:3); phosphatidylcholines (PC) such as PC(36:4) and PC(34:3); cardiolipins (CL) including CL(72:8) and CL(72:9); and lysophosphatidyl ethanolamine (LPE 18:0) (Fig. 3E). In addition, oxygenated fatty acids (OxFA) under the fatty acyls (FA) category were identified, such as FA18:2;O2 (Figure 5A) and FA 20:0;O. The classification performance of these lipid biomarkers was further assessed using receiver operating characteristic (ROC) curves. The area under the ROC curve (AUC) was used to assess model performance (Fig. 3F). Both model specificity and sensitivity were confirmed using ROC curves, which efficiently distinguished rotenone-exposed *Drosophila* from controls.

**Table 1:**
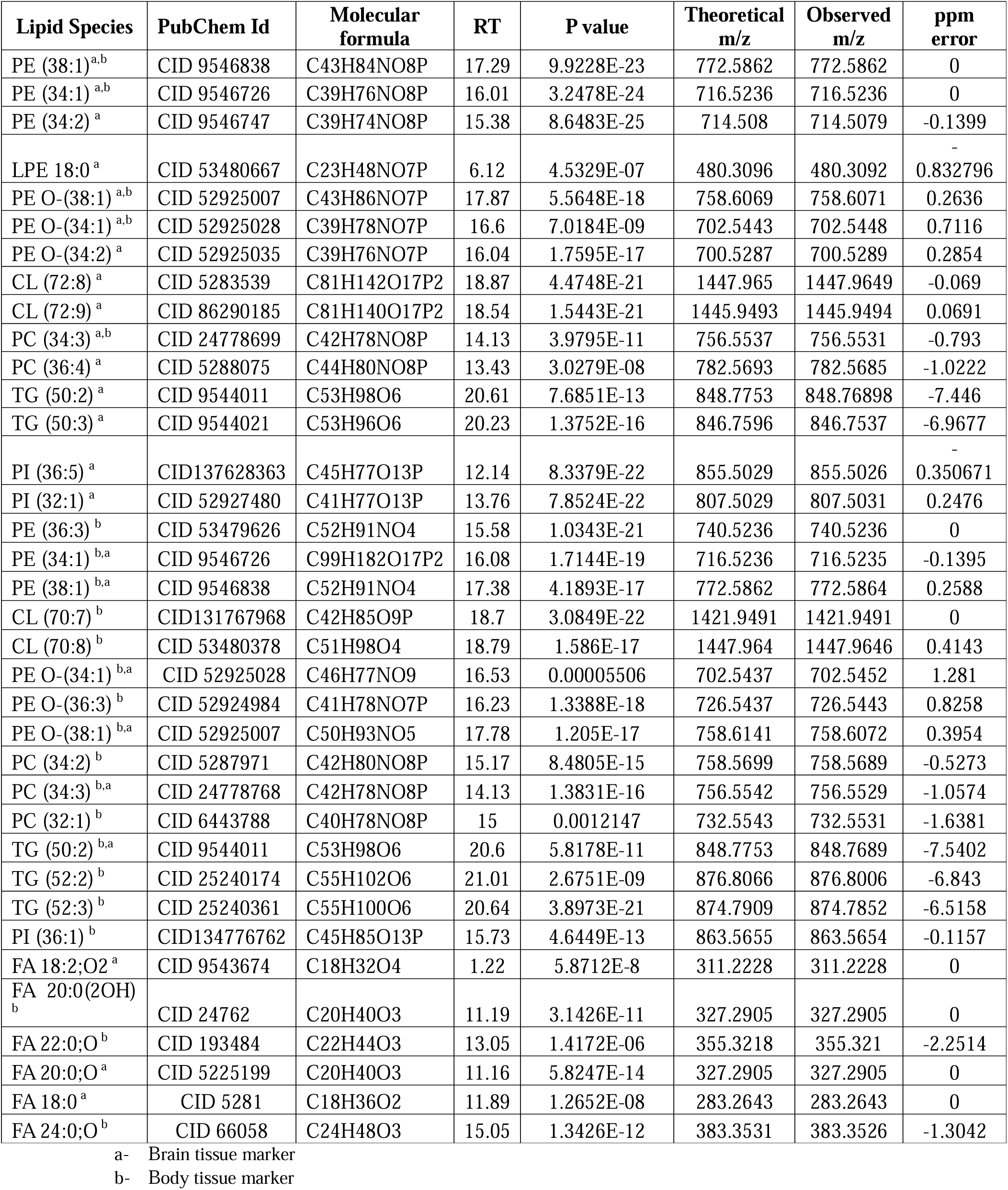
Significantly altered lipid markers (VIP >1) for rotenone exposure in brain and body tissues.

#### 3.3.3. Chronic rotenone exposure alters *Drosophila* body lipidome

Furthermore, LC-HRAMS-based metabolite profiling was performed to comprehensively identify the altered lipid profiles of body tissue after *Drosophila* exposure to rotenone. LC-MS metabolite profiling using ESI (+) and ESI (-) modes identified 321 metabolites. Multivariate analysis was performed to analyze untargeted LC-MS-based lipid profiling. For the ESI (-) ionisation mode, the first two main components of the PCA model accounted for 92.1% of the total variation, with PC1 explaining 83.2% and PC2 explaining 8.9% (Fig. 4A). Supplementary In the case of ESI (+) ionisation mode, the first two main components of the PCA model accounted for 71% of total variation, with PC1 explaining 44.7% and PC2 explaining 26.3% (Fig. 4B). Figures S7A and S7B shows the cumulative explained variances for the PCs. The supervised PLS-DA model was applied to all fly samples exposed to different concentrations of rotenone. Component 1 explained 82.9% of the variation between control, 125 µM treated, and 250 µM, 500 µM treated samples, while component 2 explained 9.2% of the variation among sample groups (Fig. S7C) for ESI (-) mode. In the ESI (+) mode, the first two components accounted 66.7% of the variation, with the first component accounting for 42.7% and the second for 24% of the variation in dose-dependent patterns among sample groups (Fig. S7D). The polynomial model’s predictive accuracy and fit were determined using five-fold cross-validation. The PLS-DA model’s accuracy of 0.9721, R² = 0.94444, and Q² = 0.91066, indicate a strong fit (Figs. 4C & S8A). To validate the supervised models, 1,000 permutation tests were performed. The study of these distributions revealed that the model’s ability to reliably predict sample group profiles was very significant, with a P-value of less than 0.001 (see Supplementary Fig. S8B and C). To compare control and rotenone-treated body sample extracts, Pearson correlation was used, showing a strong positive correlation between each sample group and its corresponding control and treatment group (Fig. 4D). Variables in important projection identified significant metabolites altered in response to chronic rotenone exposure (Table 1). These include PE class lipids, PE lipids such as PE(38:1), PE(36:3), PE(34:1), and PE O-lipids PE O-(38:1), PE O-(36:3), PE O-(34:1) and TG lipids TG(50:2), TG(52:2), TG(52:3) and PC lipids PC(34:2), PC(34:3), PC(32:1) and cardiolipins such as CL(70:7), CL(72:8) and PI lipids PI(36:1) (**Fig. 4E**). The AUC was used to assess model performance (Fig. 4F). Both model specificity and sensitivity were confirmed using ROC curves, which efficiently distinguished rotenone-exposed *Drosophila* from controls. In addition to these, we found some oxygenated fatty acids including FA 20:0(20H), FA 22:0;0, FA 24:0; O (Fig. 5)

**Figure 4:**
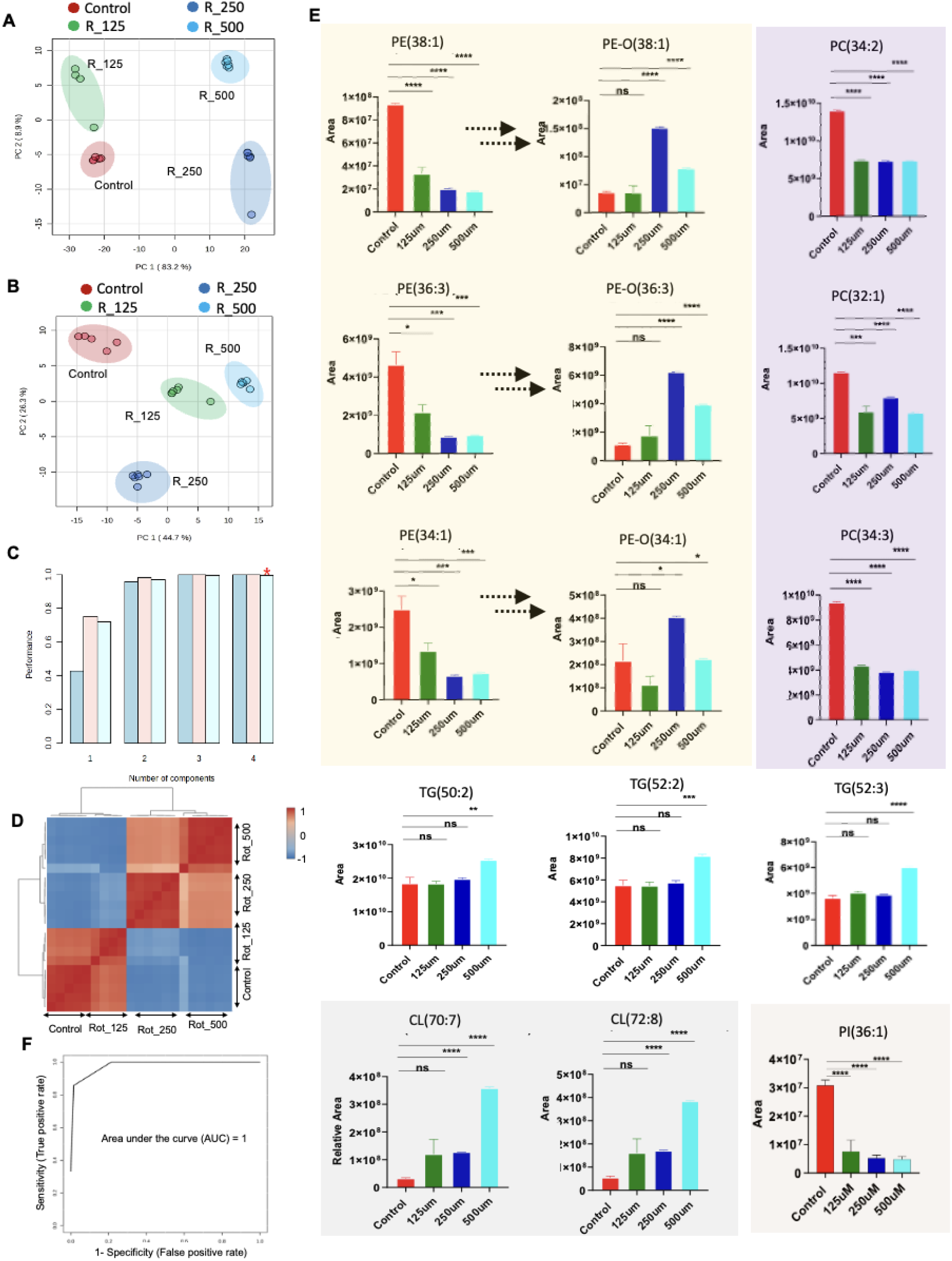
Significantly altered marker lipids in body tissue under rotenone exposure. A. For the ESI (-) ionisation mode, the first two main components of the PCA model accounted for 92.1% of the total variation, with PC1 explaining 83.2% and PC2 explaining 8.9% B. ESI (+) ionisation mode, the first two main components of the PCA model accounted for 71% of total variation, with PC1 explaining 44.7% and PC2 explaining 26.3% C. The PLS-DA model’s accuracy of 0.9721, R² = 0.94444, and Q² = 0.91066, indicate a strong fit D. Pearson correlation E. Marker lipids including PE(38:1), PE(36:3), PE(34:1), and PE O-lipids PE O-(38:1), PE O-(36:3), PE O-(34:1) and TG(50:2), TG(52:2), TG(52:3) and PC(34:2), PC(34:3), PC(32:1) and CL(70:7), CL(72:8) and PI(36:1) F. Area under the ROC curve (AUC) was used to assess model performance that distinguished rotenone-exposed *Drosophila* from controls.

**Figure 5:**
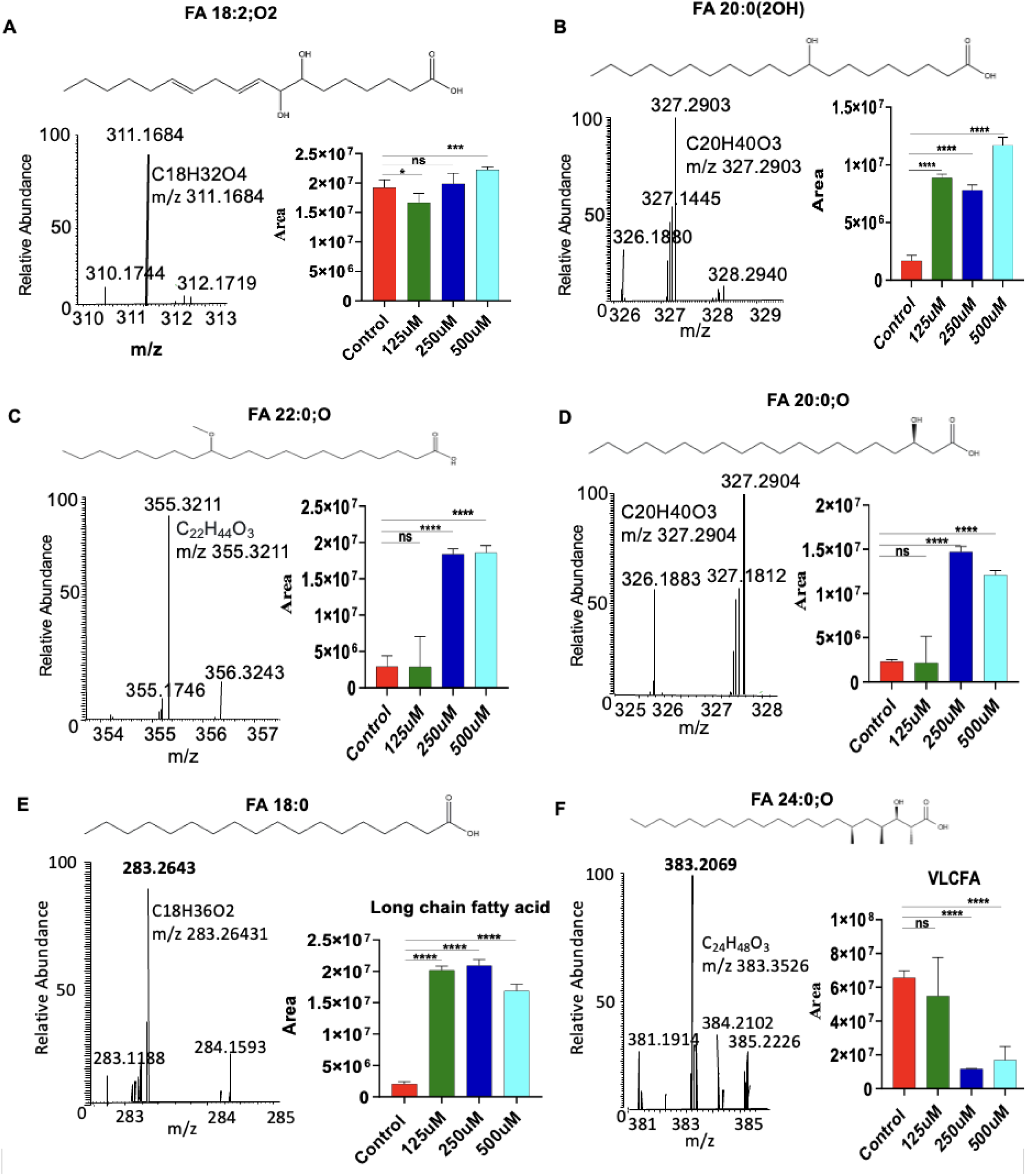
Identified of oxygenated fatty acyl (OxFA) lipids in both brain and body tissue of *D. melanogaster* under rotenone toxicity. A. FA 18:2;O2 in brain tissue B. FA 20:0(2OH) in body tissue C. FA 22:0;O in body tissue D. FA 20:0;O in brain tissue E. Long chain fatty acid (FA 18:0) in brain tissue. F. Oxygenated very long chain fatty acid (FA 24:0; O) in body tissue.

### 3.4 Chronic-exposure to rotenone elevates ROS levels

Following the observation of decreased behavioral activity, increased mortality, and disrupted lipid metabolism after chronic rotenone exposure, we investigated oxidative stress markers to assess cellular damage in the brains of *D. melanogaster* (Fig. 6A-D). ROS levels were quantified. Notably, ROS fluorescence intensity, measured via DCF-DA was significantly elevated in the brain of *D. melanogaster* treated with 250 µM and 500 µM rotenone (Fig. 6E), (*P* < 0.0001 for both concentrations), whereas the 125 µM group showed no statistically significant difference compared to controls. This result was further validated with DCF-DA staining and confocal imaging with the scale bar 50 µm and a 10X image of the brain. In contrast, in the body tissues, ROS production was significantly higher in the (Fig. 6F), 125 µM (*P* = 0.0002) and 250 µM and 500uM (*P*= 0.0018) rotenone-treated groups as compared to controls.

**Figure 6:**
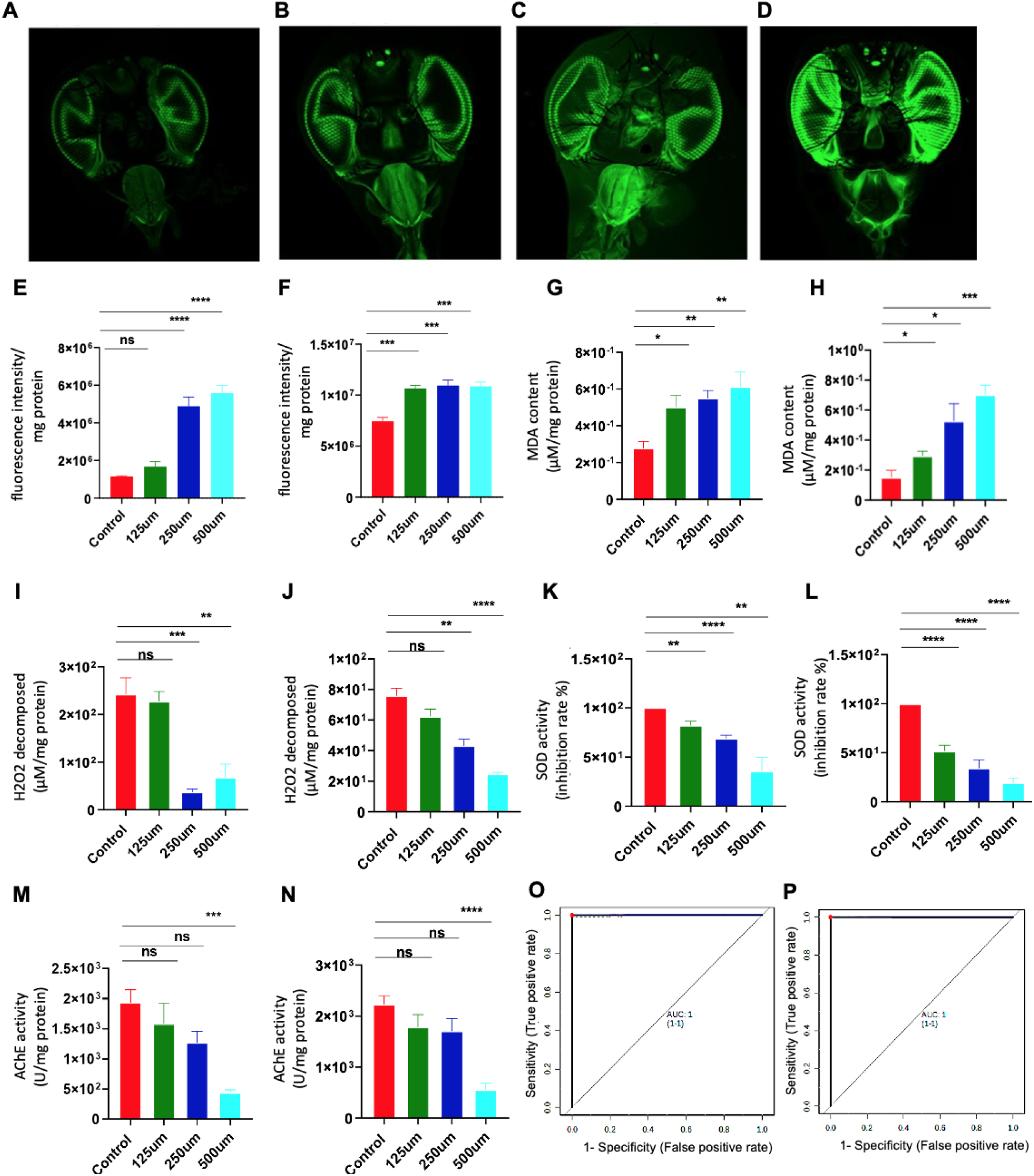
Validation assay for confirmation of rotenone effects. **A**. ROS intensity confocal image of Control brain tissue B. ROS intensity confocal image of 125uM rotenone treated brain tissue C. ROS intensity confocal image of 250uM rotenone treated brain tissue D. ROS intensity confocal image of 500uM rotenone treated brain tissue E & F. Reactive oxygen species (ROS) levels in brain and body tissue G & H. Malondialdehyde (MDA) levels in brain and body tissue I & J. Catalase or H2O2 decomposition activity in brain and body tissue K & L. SOD activity in brain and body tissue **M &N**. AChE activity in brain and body tissue assay in samples of the *D. melanogaster* exposed to Rotenone O & P. Area under the ROC curve (AUC) model performance of these biochemical parameters. Values are expressed as mean (s) ± standard error of the mean (SEM). *Significant difference compared to the control group (*p*\* < 0.05, *p*\*\*<0.005, *p*\*\*\*\*<0.0001 Student’s t-test).

### 3.5 Chronic rotenone exposure increases MDA production

To further clarify the increase in ROS levels observed in the induced model, we measured lipid peroxidation using the MDA assay, a widely recognized biomarker of oxidative stress in biological tissues. MDA is a secondary product formed during the oxidation of polyunsaturated fatty acids (PUFA) and serves as a reliable indicator of lipid peroxidation [40], reflecting the extent of oxidative damage to cellular membranes. Consistent with the elevated ROS levels, MDA concentrations were significantly higher in the brain of rotenone-treated *Drosophila* compared to controls, indicating increased lipid peroxidation (**Fig. 6G)** with 125 µM, (*P* = 0.0143), 250 µM (*P* = 0.0012) and 500 µM (*P* = 0.0052). This suggests that rotenone-induced oxidative stress resulted in substantial damage to membrane lipids in neuronal tissues. A similar pattern was observed in the body tissues, where MDA levels were significantly elevated across all rotenone-treated groups (**Fig. 6H)** 125 µM (*P* = 0.0399), 250 µM, (*P*= 0.0182) and 500 µM (*P*= 0.0002), further confirming the systemic nature of lipid peroxidation.

### 3.6 Chronic exposure of rotenone affects catalase activity

Under the same exposure conditions, we assessed the activity of key detoxifying enzymes, particularly catalase, which plays a pivotal role in the antioxidant defence system. Catalase is an essential enzyme that mitigates oxidative stress by catalyzing the decomposition of H□O□ into water and oxygen. This process is crucial because H□O□ is a ROS that can lead to oxidative damage, including lipid peroxidation, if not effectively neutralized. By reducing cellular H□O□ levels, catalase helps to prevent the oxidation of PUFA in cell membranes, thereby protecting cells from lipid peroxidation and overall oxidative stress. Catalase is also linked to lipid metabolism because it operates within peroxisomes, organelles involved in lipid breakdown. The reduction of catalase activity may impair the ability of peroxisomes and mitochondria to control oxidative damage, thus promoting the accumulation of oxidized lipids and other macromolecules [41]. In our study, catalase activity was significantly reduced in the brain of *Drosophila* treated with 250 µM and 500 µM rotenone (**Fig. 6I)**, 250 µM (*P* = 0.0004), and 500 µM (*P* = 0.0045). A similar reduction of catalase activity was observed in body tissues (**Fig. 6J**; 250 µM, (*P* = 0.0011) and 500 µM (*P* < 0.0001).

### 3.7 Chronic exposure of rotenone affects the SOD activity

We also assessed the relationship between altered lipid metabolites and oxidative stress by measuring the activity of SOD, another critical antioxidant enzyme. SOD is responsible for converting superoxide radicals (O□□), a highly reactive and damaging form of ROS, into less harmful molecules like H□O□, which is then further broken down by catalase. By neutralizing superoxide radicals, SOD plays a crucial role in protecting cells from oxidative damage, including lipid peroxidation [42]. Our results showed a significant decrease in SOD activity in both the brain and bodies of *D. melanogaster* exposed to rotenone, across all concentrations (**Fig. 6K**). In the brain, SOD activity was significantly reduced at 125 µM (*P* = 0.0048), 250 µM (*P*< 0.0001), and 500 µM (*P* = 0.0022). Similarly, in the body tissues, SOD activity decreased with (**Fig. 6L**) 125 µM (*P*< 0.0001), 250 µM (*P* < 0.0001), and 500 µM (*P* < 0.0001). Moreover, the lipid classes are positively correlated with SOD activity.

### 3.8 Chronic exposure of rotenone affects AChE enzyme activity

Furthermore, we assessed the AChE activity, an enzyme required for the breakdown of the neurotransmitter acetylcholine at synaptic junctions, which is critical for maintaining adequate neuronal function and signal transmission. A reduction in AChE activity can lead to an accumulation of acetylcholine, disrupting synaptic signaling and contributing to neurotoxicity and neurodegenerative processes. Our result shows the significant reduction of AChE in brain of 500uM (*P* = 0.0002) exposed *Drosophila* (**Fig. 6M)** and with a similar reduction observed in body tissues (*P* < 0.0001) (**Fig. 6N).** The decrease in AChE activity indicates that rotenone exposure impairs cholinergic neurotransmission, potentially leading to synaptic dysfunction and contributing to the neurodegenerative effects observed in this model. These findings suggest that rotenone exposure induces considerable oxidative stress, as seen by increased lipid peroxidation and reduced activity of key antioxidant enzymes such as catalase and SOD. The impairment of both AChE and these antioxidant defence mechanisms further exacerbates the cellular damage and neurodegeneration associated with exposure of rotenone, highlighting the multifaceted effect of oxidative stress on neuronal function.

The AUC was used to assess model performance (Fig. 6O,P) of these biochemical parameters and found that the model significantly distinguished rotenone-exposed *Drosophila* from controls (Figs 7 O & P, S9A-C).

**Figure 7:**
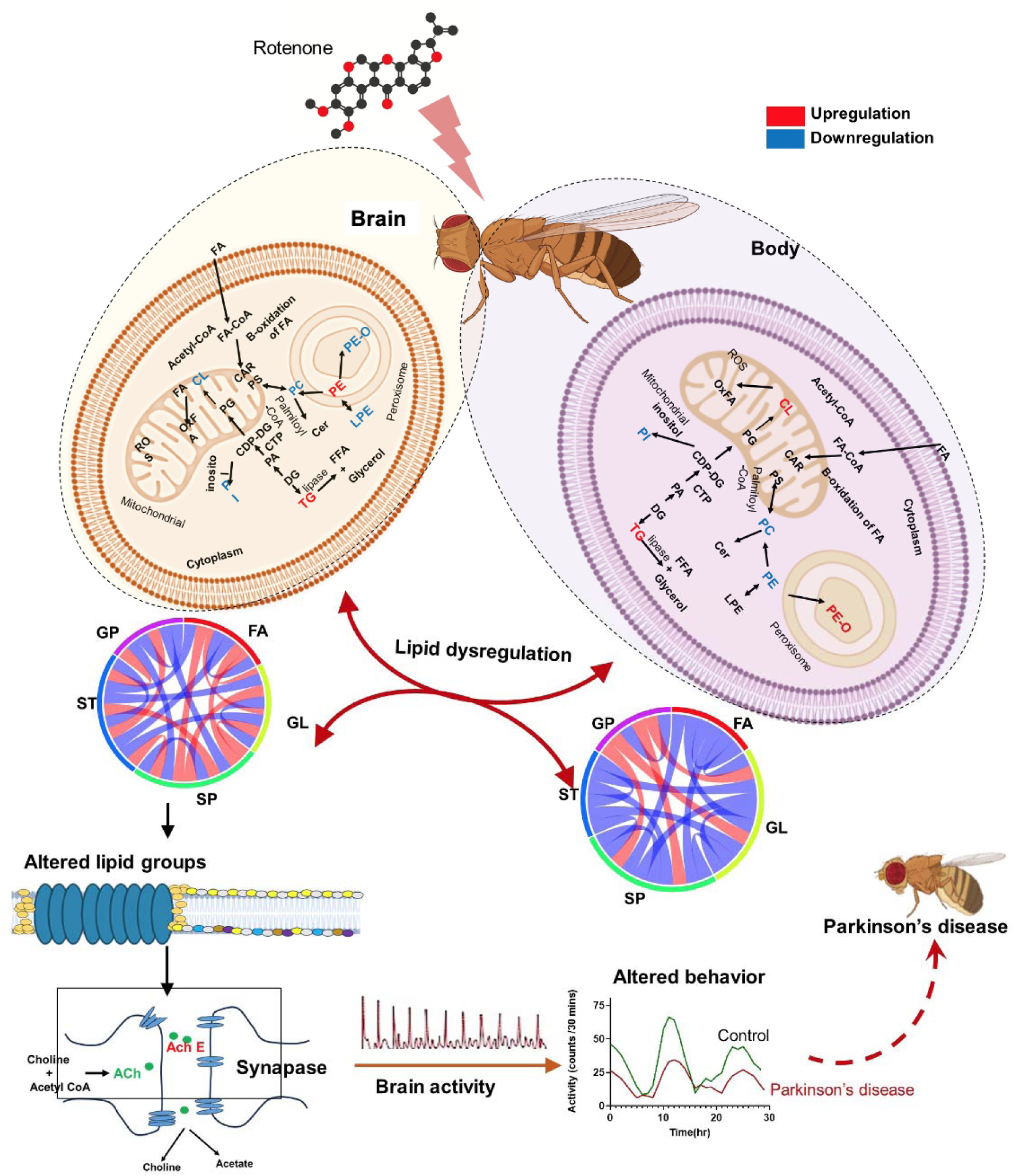
Schematic of pathway localization of altered lipids in brain and body tissue of *D. melanogaster* under chronic exposure of rotenone. Free fatty acids (FFA), Phosphatidic acid (PA), Acylcarnitine (CAR), Diacylglycerol (DG), Triacylglycerol (TG), Cardiolipin (CL), Lysophosphatidylethanolamine (LPE), Phosphatidylcholine (PC), Phosphatidylethanolamine (PE), Phosphatidylglycerol (PG), Phosphatidylinositol (PI), Phosphatidylserine (PS), Ceramides (CER) Ether-linked Phosphatidylethanolamine (PE O-).

## 4. Discussion

The continuous exposure of humans to a wide variety of environmental toxicants raises critical environmental and health concerns, as these substances can cause behavioral and cellular changes that lead to the onset of numerous disorders, including neurodegenerative diseases. Exposure to biopesticide rotenone induces PD pathogenesis, possibly mediating through disruption of mitochondrial function by inhibiting complex I, leading to increased oxidative stress which could damage cell membranes and lipid homeostasis within the dopaminergic neurons. In this study, we investigated the effects of chronic rotenone exposure on behavior, as well as on brain and peripheral lipid metabolism in *Drosophila melanogaster*. The results show that chronic exposure to rotenone induces PD-like phenotypes coupled with reduced brain tyrosine hydroxylase (TH) activity and dopamine levels. Our high-throughput untargeted lipid profiling indicates that rotenone exposure alters brain and peripheral lipidomes differently. Alteration of neurochemical and oxidative stress markers pointing toward rotenone-induced dyslipidemia could disrupt the neurotransmission and antioxidant status of cells in the progression of PD.

Rotenone has been used to induce sporadic PD in *Drosophila melanogaster* [43,44]. Here, we investigated how chronic exposure to rotenone affects locomotion and daily rhythms of *Drosophila melanogaster*. We reported that rotenone-laced diets cause a significant drop-in total activity and impaired climbing abilities in the negative geotaxis test in a dose-dependent manner. Rotenone treatments also affect circadian rhythms and diurnal activities in a dose-dependent manner. The flies displayed reduced activity levels and delayed response before the dark phase, accompanied by significant phase shifts in circadian activity. These disruptions align with previous findings in PD patients, who experience alterations in daily activity rhythms that include reduced amplitude, decreased diurnal activity, and diminished nocturnal rest [45,46]. Further, rotenone-exposed flies show reduced brain dopamine levels, and TH activity, a key enzyme in dopamine synthesis, establishes a robust PD-like phenotype both neurochemically and behaviourally.

To understand how chronic rotenone exposure, which induces behavior deficiency and PD, affects lipid metabolism, we surveyed global lipidomes of fly brain and bodies. Our analysis reported a unique pattern of PE lipids and their ether linked PE lipids (PE O-) in the brain and peripheral body tissues. Rotenone significantly reduces PE O-lipids and increases PE lipids in brain tissue; intriguingly, this trend is reversed in peripheral body tissues. PE O-lipids are significant structural constituents of the cell membranes in the brain cells that ensure structural integrity and protection from oxidative stress due to their vinyl ether bond which is less susceptible to peroxidation than regular ester-linked lipids [47]. We reported a low level of PE O-lipids like PE O-(38:1), PE O-(34:1), PE O-(34:2) in the brain (Fig. 3E) of the fly treated with rotenone. A similar dip in PE O-levels was reported previously under neurotoxicant exposure [47]. As PE lipids serve as precursors for PE O-synthesis through the peroxisomal-dependent multistep pathway, disruptions in peroxisomal function likely contribute to the observed lipidomic changes [47]. Rotenone-induced oxidative stress could compromise peroxisomal enzymes crucial for PE O-biosynthesis, such as glyceronephosphate O-acyltransferase (GNPAT) and alkylglyceronephosphate synthase (AGPS). Of note, these enzymes are highly sensitive to oxidative damage, which could impair their ability to maintain PE O-lipids. Contrary to the brain, body PE O-lipids such as PE O-(38:1), PE O-(36:3), PE O-(34:1) level was significantly increased in rotenone-exposed flies (Fig.4E). This rise could explain a protective compensatory mechanism where the body attempts to counteract the widespread oxidative stress induced by rotenone. Elevated PE O-levels in body tissue may serve as a buffer to protect cellular components, particularly highly vascularized tissues, from rotenone-induced oxidative damage. This increase may also suggest the redirection of metabolic resources away from the central nervous system to peripheral tissues, possibly leading to a net depletion of these essential lipids in the brain.

We observed surprising fluctuation in PE levels both in the brain (increase) and body (decrease) in response to rotenone treatments, despite it being the precursor of PE O-lipids. Elevated levels of PE in the substantia nigra are also reported in PD patients post-mortem brain samples [48]. PE lipids play a crucial role in maintaining membrane fluidity and curvature, properties essential for synaptic vesicle fusion, and neurotransmitter release in the brain. The accumulation of PE lipids in the brain may be due to the peroxisomal enzyme oxidation that leads to the course of non-occurring forward reaction of PE to their plasmalogen PE O-lipids. Additionally, PE is synthesized through both the Kennedy pathway (CDP-Ethanolamine pathway) and the decarboxylation of phosphatidylserine (PS) in mitochondria, suggesting that rotenone exposure may activate these pathways as a response to increased demand for structural lipids to sustain membrane stability [48]. Further, this accumulation also indicates an adaptive response to bolster plasmalogen synthesis, potentially indicating an active attempt by neural cells to replenish PE O-lipids levels that have been depleted due to rotenone exposure.

Conversely, the decrease in PE lipid precursors in peripheral tissues may occur as a trade-off to prioritize the brain’s need for these lipids. Since plasmalogen synthesis requires specific precursor molecules PE, the redirection of precursor PE lipids toward the brain could compromise the availability of these lipids in peripheral tissues, resulting in their decrease. This redistribution underscores the CNS’s critical need to maintain lipid homeostasis and protect neural tissue against oxidative damage in response to neurotoxicants, even at the expense of peripheral lipid levels. The divergence in the alteration of PE lipids and their plasmalogen form (PE O-) in the brain and body, highlights intricate, tissue-specific metabolic adaptations and protective mechanisms activated against neurotoxic stress caused by rotenone exposure. Our findings are supported by the evidence from a study conducted on PD mice that demonstrated reduced levels of PE O-lipids in the brain tissue [49].

The most significant lipids altered after PE/PE O-lipids are triglycerides (TGs). TGs are crucial lipids involved in energy storage, transport, and cellular homeostasis. In the brain, they contribute indirectly to the structural integrity of neurons and glial cells, as well as to neuroprotection and neural metabolism [50]. We observed a significant elevation of TG lipids levels in both brain and peripheral body tissues following rotenone exposure. Specifically, TG (50:2), and TG (52:3) were elevated in both tissues, while TG (52:2) showed increased levels exclusively in peripheral body tissues (Fig. 3E & 4E). This increase represents a complex and adaptive metabolic response to counter oxidative cellular damage induced by rotenone. These findings align with human studies, where elevated levels of TG lipids were found in PD brain tissues [50]. Furthermore, several lipidomics investigations of biological fluids from PD patients, including plasms, serum, and sebum, have similarly revealed altered TG metabolism, which mirrors the changes observed in *Drosophila* peripheral tissues. Rotenone can cause oxidative stress and inflammation, prompting an adaptive lipid response where cells increase TG synthesis as a protective mechanism. Prolonged chronic exposure can lead to lipid accumulation, membrane dysfunction, and mitochondrial impairment, which contribute to neurodegeneration. Furthermore, these changes in TG levels in both the brain and body tissue can influence each other through the brain-body axis as the excess TGs in the peripheral tissues can alter metabolic signaling and inflammatory mediators communicate with the brain, potentially exacerbating neuroinflammation and toxicity. Conversely, elevated TGs in the brain could influence CNS functions that regulate peripheral lipid metabolism, contributing to systemic dyslipidemia. This interdependence highlights the significance of the observed TG lipids increase in both central and peripheral systems during rotenone exposure. The increased TG levels can serve as a crucial biomarker for rotenone exposure and further, this could offer insight into potential therapeutic interventions by targeting TG lipid clearance through pharmacological or dietary interventions that help mitigate the harmful effects of rotenone.

Another lipid form altered during chronic rotenone exposure is PCs. Rotenone decreases PC lipid levels throughout the brain and body indicating widespread disruption in lipid homeostasis. PCs are a crucial class of phospholipids widely found in cell membranes and play a functional role in cell signaling, membrane dynamics, lipid transport, metabolism more importantly neuroprotection, brain health, and in cellular repair. We reported, PC lipids exhibited tissue-specific downregulation in response to rotenone exposure. In brain tissues, PC (36:4) and PC (34:3) were significantly reduced (Fig.3E), while in peripheral body tissue, a similar trend was observed for PC (34:2), PC (34:3), and PC (32:1) (Fig.4E). This reduction in PC lipids suggests impaired lipid biosynthesis, potentially driven by oxidative stress and rotenone-induced mitochondrial dysfunction. These findings are consistent with a lipidomics study performed on human PD brain samples reporting reduced levels of PC lipids in affected areas [50].

We hypothesize that a significant decrease in PCs in the brain because these lipids undergo oxidation, where fatty acid chains in the PC molecules disintegrate, altering the fluidity and integrity of neuronal membranes. This degradation of PCs leads to subsequent weakening of membrane stability, making neurons more susceptible to damage. Furthermore, PC lipids are the major source of choline, which is essential for synthesizing acetylcholine (ACh), a neurotransmitter involved in memory, learning, and muscle control. Degradation of PCs by rotenone can reduce choline availability, leading to impaired neurotransmitter synthesis and causing neurological disorders. Disruption of PC lipids in human brain samples has been linked to the development and progression of neurodegenerative diseases, including Alzheimer’s, Parkinson’s, and Huntington’s [48,50]. Similarly, the significant decline of PCs in the peripheral body tissue suggests that rotenone may interfere with the pathways responsible for the synthesis of PCs leading to metabolic dysregulation.

PI lipids are essential components of cell membranes and contribute to membrane integrity, fluidity, and dynamics were also significantly altered in *Drosophila* in response to chronic rotenone exposure. Here, we identified rotenone exposure led to a notable decrease in PI lipids such as PI (36:5), PI (32:1) in the brain tissues (Fig.3E). Again, this reduction aligns with findings from human brain studies, which reported decreased levels of PI lipids in affected regions [48]. In the brain, these PI lipids are critical for neuronal functions and the production of phosphatidylinositol 4,5-biphosphate (PIP2), which upon stimulation, are hydrolyzed to generate signaling molecules like inositol triphosphate (IP3) and diacylglycerol (DAG). These molecules play a critical role in regulating Ca^+2^ release from smooth endoplasmic reticulum and protein kinase C activation, contributing vital processes for learning, memory, and overall cognitive function [51]. Some studies have linked disruptions of PI lipid metabolism to neurodegenerative diseases [48]. In the periphery, PI is associated with cellular stress responses. In our study, we found a depletion of PI (36:1) in the fly’s body (Fig.4E) following rotenone treatment. The PI lipids in the body are involved in the regulation of lipid droplets, crucial for energy storage and mobilization, and influence the metabolism of triglycerides. Additionally, PI lipids play an important role in the cellular response to oxidative stress and inflammation, promoting survival and repair mechanisms in the face of cellular damage [51]. The dysregulated PI lipids in fly bodies may be due to pathophysiological conditions induced by rotenone exposure.

We also reported significant alterations of cardiolipins (CLs) in the brain. CL is a unique phospholipid predominantly abundant in the inner mitochondrial membrane and contributes a crucial role in mitochondrial function and overall cellular health [50]. These anionic phospholipids play important roles in peripheral tissues, such as maintaining membrane fluidity and regulating mitochondrial processes. CL is present in both neuronal and non-neuronal glial cells within the CNS, supporting mitochondrial function, and maintaining brain cell viability [51,52]. CL orchestrates intracellular and mitochondrial signaling to facilitate efficient electron transport and mitochondrial membrane fusion and fission [52]. Associated with the inner mitochondrial membrane, CL is essential for stabilizing respiratory supercomplexes formed by the aggregation of electron transport chain complexes I, III, and IV. Moreover, CL serves as an elimination signal in injured or diseased neurons, mediating the selective removal of dysfunctional mitochondria through mitophagy. Additionally, CL contributes to the regulation of programmed cell death via apoptosis [53]. Excessive ROS production from dysfunctional mitochondria can harm cellular membranes and organelles. Mitophagy is a mechanism that selectively degrades defective mitochondria to reduce the damage caused by high ROS levels. The mitochondrial isoform of creatine kinase helps move CL from the inner to outer mitochondrial membranes. By stabilizing electron transport chain (ETC) complexes, it allows for effective ATP synthesis via oxidative phosphorylation, which is critical for meeting brain tissue’s energy demands. Our study reported the downregulation of CL lipids such as CL (72:8), CL (72:9) (Fig.3E) in the brain, and upregulation of CL (70:7), CL (72:8) in the body (Fig.4E) of fly treated with rotenone. The decline of CLs in the brain is attributed to the inhibition of complex I by rotenone in mitochondria that hampers the electron transport chain, reducing ATP production and subsequently leading to increasing ROS levels. Again, this is consistent with previous findings in human PD studies [54]. Other than that, a shotgun lipidomics study on amygdala homogenates from PD brains identified a decrease in CL levels, further supporting the link between mitochondrial dysfunction and altered lipid metabolism in PD [55]. The CLs in mitochondria are highly susceptible to ROS-induced oxidative damage that degrades the existing CL molecules and impairs their synthesis. Because neurons heavily rely on mitochondrial function for energy production, reduced cardiolipin levels can severely impact brain health, contributing to neurodegenerative processes. Declining CLs levels have been linked to neurodegenerative diseases, such as Alzheimer’s and Parkinson’s, highlighting their role in maintaining neuronal health. These CLs can serve as potential biomarkers for rotenone exposure.

More importantly, in the present study, we found significantly elevated levels of oxidized fatty acids derived from fatty acyl lipids (OxFA), FA 18:2; O2, and FA 22:0; O (Fig.5) in the brain tissue of rotenone-treated flies. Furthermore, we also found increased levels of OxFA lipids, FA 20:0; O and FA 20:0 (2OH) (Fig.5) in the body tissue. The increased fatty acid oxidation levels in rotenone-exposed animal models highlight significant perturbation in cellular energy homeostasis, a hallmark of mitochondrial dysfunction. This disruption caused by ROS triggered compensatory metabolic adaptations, including increased reliance on fatty acid β-oxidation to meet energy demands supported by an in-vitro study, in which rotenone induces fatty acids β-oxidation in SH-SY5Y cells [56]. Mitochondria play a crucial role in β-oxidation, where fatty acids are broken down into acetyl-CoA, FADH2, and NADH. In rotenone-treated flies, inhibition of complex I impairs the utilization of NADH for oxidative phosphorylation. In addition to these, we observed a decrease in levels of very-long-chain fatty acid (VLCFA), FA (24:0;O) and an increase in plasmalogens in body tissues, under rotenone toxicity (Fig. 5F). The synthesis of VLCFA and plasmalogens is a function of peroxisome metabolic process. The decline in VLCFA levels are tightly linked to peroxisome dysfunction under rotenone induced stress conditions. This results in an energy deficit, which cells may attempt to offset by upregulating fatty acid oxidation pathways and declined VLFA levels. While fatty acid oxidation provides an alternative energy source under mitochondrial stress, its upregulation may have deleterious consequences due to excessive ROS generation and lipid metabolism imbalance.

While rotenone itself does not directly cause PD, its neurotoxic effects on mitochondrial function by ATP degradation and oxidative stress are significant contributors to the pathophysiology of the disease. More importantly, the low concentrations of rotenone in the brain are enough to partially inhibit complex I, but too low to significantly impair the respiration of brain mitochondria. It seems to be a bioenergetics effect with ATP depletion could not explain neurodegeneration. Studies have shown that oxidative damage might have been involved in this process [57]. The altered oxidative stress might have perturbed significant lipid classes through the oxidative damage. To validate this, we performed ROS assay and discovered the dose-dependent effect of rotenone on ROS production with distinct patterns between brain and body tissues. The distinct ROS patterns across tissues and concentrations offer insights into the potential for differential vulnerability and adaptive response to oxidative stress within the organism.

We found elevated level of ROS in *Drosophila* similar to the PD human post-mortem brain [58]. The oxidative stress, resulting from ROS, can lead to lipid peroxidation, producing critical by-product MDA, a highly reactive aldehyde that has crosslinks with proteins and nucleic acids, further exacerbating cellular function through damages cellular membranes, disrupting membrane fluidity. Considering this, we next performed the MDA assay and found a significant elevation in MDA levels in both brain and body tissues which aligns with the excessive ROS production, reinforcing the role of oxidative stress in rotenone-induced cellular damage. Human PD patients have a high MDA level in the brain. This lipid peroxidation is particularly detrimental in neuronal cells, where it contributes to membrane instability, mitochondrial dysfunction, and the progressive degeneration of dopaminergic neurons, characteristic of PD. Thus, the increase in MDA levels underscores the critical link between oxidative stress, lipid peroxidation, and neurodegeneration in our rotenone-exposed *Drosophila* model.

The imbalance in ROS production can directly affect catalase activity as it is an essential antioxidant enzyme responsible for decomposing hydrogen peroxide. We observed reduced catalase activity in both brain and body tissues after their exposure to rotenone. Under high oxidative stress, excessive ROS can overwhelm catalase activity, leading to reduced enzymatic efficiency and impaired detoxification. This reduction in catalase activity suggests that the rotenone-treated flies are less equipped to neutralize hydrogen peroxide, thereby exacerbating oxidative stress within cells. These findings align with human studies reporting deficits in catalase activity in postmortem brain tissues of PD patients, particularly in the substantia nigra and putamen regions [59]. Additionally, a marked reduction in catalase activity has been observed in the erythrocytes of PD patients compared to healthy controls [60]. This diminished catalase activity likely contributes to the increased lipid peroxidation noted in these tissues, as reduced catalase availability permits the accumulation of ROS, thereby promoting lipid oxidation. Consequently, the decrease in catalase activity exacerbates oxidative damage and contributes to the progression of cellular dysfunction, particularly in neuronal tissues where lipid peroxidation can lead to neurodegeneration.

Oxidative stress, by altering catalase activity and ROS dynamics, may influence SOD activity, a critical enzyme that catalyzes the conversion of superoxide radicals into hydrogen peroxide, which is subsequently degraded by catalase. Considering this, we measured catalase activity and found decreased enzyme activity in both brain and body tissues. It may be attributed to that under chronic rotenone exposure conditions, SOD can become overwhelmed due to oxidative modification to its protein structure. Impaired SOD function may result in the accumulation of superoxide radicals, causing cellular oxidative damage. This decrease in SOD activity indicates a diminished capacity to neutralize superoxide radicals, leading to an accumulation of ROS and enhanced oxidative stress SOD enzyme a common biomarker for Human PD patient’s brains was decreased. As a result, the increased oxidative stress likely exacerbates lipid peroxidation and other forms of cellular damage.

The enzyme AChE breaks down ACh into choline and acetate and a neurotransmitter, which is synthesized from choline and acetyl-CoA. The choline component of ACh derives from PC lipids, and the metabolism of PC is indirectly linked to Ach levels and, consequently, to AChE activity. Reduction in PC lipids in rotenone-exposed fly tissues was observed in the present study and may directly affect AChE activity. Considering this, AChE was measured in *Drosophila* brain and body tissues and found that its levels were decreased in both tissues after rotenone exposure similar to the previously reported human PD brain study where AChE activity was significantly decreased in the medial occipital cortex [61]. The decrease in ACheE activity indicates that rotenone exposure impairs cholinergic neurotransmission, potentially leading to synaptic dysfunction and contributing to the neurodegenerative effects observed in this model. It indicates PC lipids and AChE are closely related in the context of maintaining membrane dynamics after their exposure to rotenone. Schematic of altered lipid biosynthesis pathway under chronic exposure of rotenone showing the significant regulation of individual class of lipids in brain and body tissue (Fig.7).

More importantly, the present study revealed significant changes in lipid profiles, including lipid metabolites that correlated with those observed in postmortem brain samples from human PD subjects (Table 2). This cross-species correlation underscores translations relevance of *Drosophila* model in studying PD. By bridging the gap between experimental models and human pathology, this work, hold promise for advancing clinical interventions and enhancing therapeutic strategies for PD.

**Table 2:**
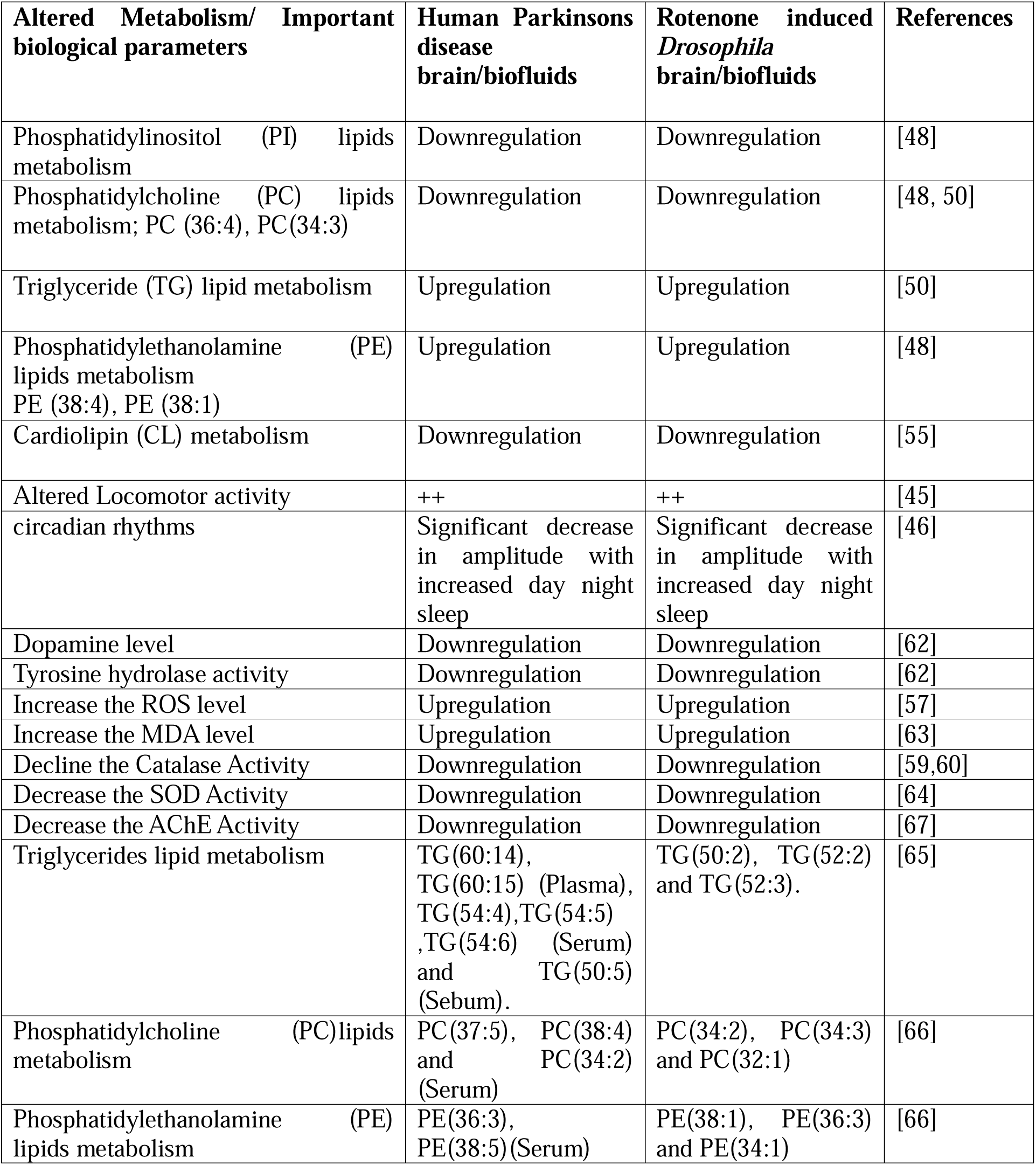
Correlation of altered lipid metabolites with human samples.

## 5. Conclusion

Current research suggests that chronic exposure to the environmental toxin rotenone poses a threat by contributing to neurotoxicity and the pathogenesis PD. This toxicity is mediated through the alterations of systemic lipid compositions. Comprehensive lipid profiling reveals that rotenone exposure differentially alters global lipidomes of the brain and peripheral tissues of *Drosophila melanogaster*. The lipid biomarkers from PE, TG, PC, CL, and PI lipid classes were significantly perturbed. Notably, these lipids play critical roles in membrane formation, cellular organelle maintenance, signal transduction, and energy storage, essential for survival. The depletion of the brain PE O-lipids and concomitant rise of PE, whereas augmentation of peripheral PE O-lipids and concomitant fall of PE suggest tissue-specific metabolic adaptations and protective mechanisms to mitigate oxidative stress caused by neurotoxicant exposure. Moreover, increased levels of oxidized fatty acids in both brain and peripheral tissues are indicative of disruption in cellular energy balance, driven by mitochondrial dysfunction. During rotenone exposure, biochemical parameters such as dopamine, TH, catalase, MDA, ROS, antioxidant enzyme SOD, and neurochemical AChE change, indicating changes in the antioxidant status and activity of important neurotransmission enzymes. In sum, these findings highlight the adaptive lipid changes at both intra-tissue and inter-tissue levels in response to rotenone toxicity. At the same time, these lipids could serve as potential biomarkers to manage and treat PD pathogenesis.

### Environmental implication

The environmental implications of this study are profound, as they underscore the potential risks associated with chronic rotenone exposure, which is commonly used in agriculture. The findings indicate that even low, environmentally relevant concentrations of rotenone can lead to significant disruptions in lipid metabolism, particularly in the brain. This disruption affects mitochondrial and peroxisomal functions, contributes to oxidative stress, and depletes crucial dopaminergic neurons, all of which are central to the development of neurodegenerative diseases like Parkinson’s. Furthermore, the study highlights metabolic trade-offs where the brain prioritizes neural integrity at the expense of peripheral lipid homeostasis, emphasizing the broader environmental risks of such exposure. Beyond the individual health impacts, these lipidome alterations and redox-driven metabolic shifts pose a threat to ecosystem health, particularly in regions with frequent rotenone use. As neurodegenerative disorders become more prevalent, understanding the environmental role of pesticides in brain health is critical for shaping public health policies, pesticide regulations, and preventive measures in agricultural practices, while also guiding future environmental policies to minimize exposure risks.

### CRediT authorship contribution statement

Ratnasekhar CH: Conceptualization, Formal analysis, Funding acquisition, Investigation, Project administration, Software, Supervision, Writing – original draft, – review and editing. Ashutosh Kumar Tiwari: Methodology, Data curation, Formal analysis, Validation, Visualization, Writing – review & editing. Priya Rathor: Methodology, Data curation, Formal analysis, Validation, Visualization, Writing – review & editing. Rajendra Patel: Data curation, Formal analysis, Visualization, Writing – review & editing. Nick Birse: Resources, Methodology, Writing – review & editing.

## Acknowledgements

Ratnasekhar CH acknowledges research grants from DST-Science and Engineering Research Board (SRG/2021/000750-G) and DBT (BT-RLF-Reentry-21-2020) for the Ramalingaswami re-entry fellowship programme. We are also grateful for access to the High-throughput instrumentation facility at CSIR-CIMAP. Ashutosh Kumar Tiwari was supported by DBT Fellowship. Priya Rathor was supported by UGC-JRF fellowship. We also thank Director, CIMAP for providing facility and infrastructure. Nick Birse acknowledges IGFS, QUB.

## Supplementary Data

Supplementary information consisting of Fig.S1-Fig S9 is provided.

## Data availability

Lipidomics data for can be accessed on the Metabolights platform; Study submission code: MTBLS1285 and link https://www.ebi.ac.uk/metabolights/editor/MTBLS11699/assays

## References

(1) Reedijk, J.; Dalcanale, E.; Krebs, B.; Marquardt, R.; Morbidelli, M.; Nakai, H.; Panza, L.; Poole, C.; Quack, M.; Wandelt, K. Reference Module in Chemistry, Molecular Sciences and Chemical Engineering; 2013.

(2) Innos, J.; Hickey, M. Using Rotenone to Model PD in Mice: A Review of the Role of Pharmacokinetics. Chem. Res. Toxicol. 2021, 34. 10.1021/acs.chemrestox.0c00522.

(3) Xu, L.; Wu, Z.; Li, J.; Xu, Y.; Zhou, F.; Zhang, F.; Li, D.; Zhou, L.; Liu, R. The Low-Lethal Concentrations of Rotenone and Pyrethrins Suppress the Population Growth of Rhopalosiphum Padi. Sci. Rep. 2024, 14 (1), 16570. 10.1038/s41598-024-67286-1.

(4) Betarbet, R.; Sherer, T. B.; MacKenzie, G.; Garcia-Osuna, M.; Panov, A. V; Greenamyre, J. T. Chronic Systemic Pesticide Exposure Reproduces Features of PD. Nat. Neurosci. 2000, 3 (12), 1301–1306. 10.1038/81834.

(5) Kurker, V.; Habtemichael, A.; Pittman, J.; Shelat, S.; Assessor, R.; Jones, R. D.; Jones, R.; Leader, T.; Friedman, D.; Chief, B. Ll ∼ – –. 2014, 2006.

(6) Dalu, T.; Wasserman, R. J.; Jordaan, M.; Froneman, W. P.; Weyl, O. L. F. An Assessment of the Effect of Rotenone on Selected Non-Target Aquatic Fauna. PLoS One 2015, 10 (11), e0142140. 10.1371/journal.pone.0142140.

(7) Cavoski, I.; Caboni, P.; Sarais, G.; Miano, T. Degradation and Persistence of Rotenone in Soils and Influence of Temperature Variations. J. Agric. Food Chem. 2008, 56 (17), 8066–8073. 10.1021/jf801461h.

(8) Gandara, L.; Jacoby, R.; Laurent, F.; Spatuzzi, M.; Vlachopoulos, N.; Borst, N. O.; Ekmen, G.; Potel, C. M.; Garrido-Rodriguez, M.; Böhmert, A. L.; Misunou, N.; Bartmanski, B. J.; Li, X. C.; Kutra, D.; Hériché, J.-K.; Tischer, C.; Zimmermann-Kogadeeva, M.; Ingham, V. A.; Savitski, M. M.; Masson, J.-B.; Zimmermann, M.; Crocker, J. Pervasive Sublethal Effects of Agrochemicals on Insects at Environmentally Relevant Concentrations. Science 2024, 386 (6720), 446–453. 10.1126/science.ado0251.

(9) Causes, W. Understanding_Parkinson_s_dise.PDF. 2020, 1–2.

(10) Tanner, C. M.; Kamel, F.; Ross, G. W.; Hoppin, J. A.; Goldman, S. M.; Korell, M.; Marras, C.; Bhudhikanok, G. S.; Kasten, M.; Chade, A. R.; Comyns, K.; Richards, M. B.; Meng, C.; Priestley, B.; Fernandez, H. H.; Cambi, F.; Umbach, D. M.; Blair, A.; Sandler, D. P.; Langston, J. W. Rotenone, Paraquat, and PD. Environ. Health Perspect. 2011, 119 (6), 866–872. 10.1289/ehp.1002839.

(11) Pouchieu, C.; Piel, C.; Carles, C.; Gruber, A.; Helmer, C.; Tual, S.; Marcotullio, E.; Lebailly, P.; Baldi, I. Pesticide Use in Agriculture and PD in the AGRICAN Cohort Study. Int. J. Epidemiol. 2018, 47 (1), 299–310. 10.1093/ije/dyx225.

(12) Balestrino, R.; Schapira, A. H. V. Parkinson Disease. Eur. J. Neurol. 2020, 27 (1), 27– 42. 10.1111/ene.14108.

(13) Ibarra-Gutiérrez, M. T.; Serrano-García, N.; Orozco-Ibarra, M. Rotenone-Induced Model of PD: Beyond Mitochondrial Complex I Inhibition. Mol. Neurobiol. 2023, 60 (4), 1929–1948. 10.1007/s12035-022-03193-8.

(14) Van Laar, A. D.; Webb, K. R.; Keeney, M. T.; Van Laar, V. S.; Zharikov, A.; Burton, E. A.; Hastings, T. G.; Glajch, K. E.; Hirst, W. D.; Greenamyre, J. T.; Rocha, E. M. Transient Exposure to Rotenone Causes Degeneration and Progressive Parkinsonian Motor Deficits, Neuroinflammation, and Synucleinopathy. npj Park. Dis. 2023, 9 (1), 121. 10.1038/s41531-023-00561-6.

(15) Ranasinghe, T.; Seo, Y.; Park, H.-C.; Choe, S.-K.; Cha, S.-H. Rotenone Exposure Causes Features of Parkinson’s Disease Pathology Linked with Muscle Atrophy in Developing Zebrafish Embryo. J. Hazard. Mater. 2024, 480, 136215. 10.1016/j.jhazmat.2024.136215.

(16) Cannon, J. R.; Tapias, V.; Na, H. M.; Honick, A. S.; Drolet, R. E.; Greenamyre, J. T. A Highly Reproducible Rotenone Model of PD. Neurobiol. Dis. 2009, 34 (2), 279–290. 10.1016/j.nbd.2009.01.016.

(17) Li, P.; Tian, Y.; Du, M.; Xie, Q.; Chen, Y.; Ma, L.; Huang, Y.; Yin, Z.; Xu, H.; Wu, X. Mechanism of Rotenone Toxicity against Plutella Xylostella: New Perspective from a Spatial Metabolomics and Lipidomics Study. J. Agric. Food Chem. 2023, 71 (1), 211– 222. 10.1021/acs.jafc.2c06292.

(18) Gopi, M.; Vanisree, A. J. Attenuated Levels of Phospholipids in the Striatum of Rats Infused with Rotenone Causing Hemiparkinsonism as Detected by Simple Dye-Lipid Complex. IBRO reports 2017, 3, 1–8. 10.1016/j.ibror.2017.06.001.

(19) Chen, A. Y.; Wilburn, P.; Hao, X.; Tully, T. Walking Deficits and Centrophobism in an α-Synuclein Fly Model of PD. *Genes*, Brain Behav. 2014, 13 (8), 812–820. 10.1111/gbb.12172.

(20) Shukla, A. K.; Ratnasekhar, C.; Pragya, P.; Chaouhan, H. S.; Patel, D. K.; Chowdhuri, D. K.; Mudiam, M. K. R. Metabolomic Analysis Provides Insights on Paraquat-Induced Parkinson-Like Symptoms in Drosophila Melanogaster. Mol. Neurobiol. 2016, 53 (1), 254–269. 10.1007/s12035-014-9003-3.

(21) Riabinina, O.; Potter, C. J. Drosophila: Methods and Protocols, Methods in Molecular Biology. Methods Mol. Biol. 2016, 1478, 291–302. 10.1007/978-1-4939-6371-3.

(22) Vermeer, L. M.; Higgins, C. A.; Roman, D. L.; Doorn, J. A. Real-Time Monitoring of Tyrosine Hydroxylase Activity Using a Plate Reader Assay. Anal. Biochem. 2013, 432 (1), 11–15. 10.1016/j.ab.2012.09.005.

(23) Musachio, E. A. S.; de Freitas Couto, S.; Poetini, M. R.; Bortolotto, V. C.; Dahleh, M. M. M.; Janner, D. E.; Araujo, S. M.; Ramborger, B. P.; Rohers, R.; Guerra, G. P.; Prigol, M. Bisphenol A Exposure during the Embryonic Period: Insights into Dopamine Relationship and Behavioral Disorders in Drosophila Melanogaster. Food Chem. Toxicol. 2021, 157 (August). 10.1016/j.fct.2021.112526.

(24) Caboni, P.; Sarais, G.; Vargiu, S.; Garau, V.; Ibba, A.; Cabras, P. LC–MS–MS Determination of Rotenone, Deguelin, and Rotenolone in Human Serum. Chromatographia 2008, 68, 739–745. 10.1365/s10337-008-0830-0.

(25) Guan, X. L.; Cestra, G.; Shui, G.; Kuhrs, A.; Schittenhelm, R. B.; Hafen, E.; van der Goot, F. G.; Robinett, C. C.; Gatti, M.; Gonzalez-Gaitan, M.; Wenk, M. R. Biochemical Membrane Lipidomics during Drosophila Development. Dev. Cell 2013, 24 (1), 98–111. 10.1016/j.devcel.2012.11.012.

(26) Shui, G.; Guan, X. L.; Low, C. P.; Chua, G. H.; Goh, J. S. Y.; Yang, H.; Wenk, M. R. Toward One Step Analysis of Cellular Lipidomes Using Liquid Chromatography Coupled with Mass Spectrometry: Application to Saccharomyces Cerevisiae and Schizosaccharomyces Pombe Lipidomics. Mol. Biosyst. 2010, 6 (6), 1008–1017. 10.1039/b913353d.

(27) Cajka, T.; Fiehn, O. Comprehensive Analysis of Lipids in Biological Systems by Liquid Chromatography-Mass Spectrometry. Trends Analyt. Chem. 2014, 61, 192– 206. 10.1016/j.trac.2014.04.017.

(28) Liebisch, G.; Fahy, E.; Aoki, J.; Dennis, E. A.; Durand, T.; Ejsing, C. S.; Fedorova, M.; Feussner, I.; Griffiths, W. J.; Köfeler, H.; Merrill, A. H. J.; Murphy, R. C.; O’Donnell, B.; Oskolkova, O.; Subramaniam, S.; Wakelam, M. J. O.; Spener, F. Update on LIPID MAPS Classification, Nomenclature, and Shorthand Notation for MS-Derived Lipid Structures. J. Lipid Res. 2020, 61 (12), 1539–1555. 10.1194/jlr.S120001025.

(29) Carvalho, M.; Sampaio, J. L.; Palm, W.; Brankatschk, M.; Eaton, S.; Shevchenko, A. Effects of Diet and Development on the Drosophila Lipidome. Mol. Syst. Biol. 2012, 8, 600. 10.1038/msb.2012.29.

(30) Vaccaro, A.; Kaplan Dor, Y.; Nambara, K.; Pollina, E. A.; Lin, C.; Greenberg, M. E.; Rogulja, D. Sleep Loss Can Cause Death through Accumulation of Reactive Oxygen Species in the Gut. Cell 2020, 181 (6), 1307–1328.e15. 10.1016/j.cell.2020.04.049.

(31) Pérez-Severiano, F.; Santamaría, A.; Pedraza-Chaverri, J.; Medina-Campos, O. N.; Ríos, C.; Segovia, J. Increased Formation of Reactive Oxygen Species, but No Changes in Glutathione Peroxidase Activity, in Striata of Mice Transgenic for the Huntington’s Disease Mutation. Neurochem. Res. 2004, 29 (4), 729–733. 10.1023/B:NERE.0000018843.83770.4b.

(32) Ohkawa, H.; Ohishi, N.; Yagi, K. Assay for Lipid Peroxides in Animal Tissues by Thiobarbituric Acid Reaction. Anal. Biochem. 1979, 95 (2), 351–358. 10.1016/0003-2697(79)90738-3.

(33) Anet, A.; Olakkaran, S.; Kizhakke Purayil, A.; Hunasanahally Puttaswamygowda, G. Bisphenol A Induced Oxidative Stress Mediated Genotoxicity in Drosophila Melanogaster. J. Hazard. Mater. 2019, No. March, 42–53. 10.1016/j.jhazmat.2018.07.050.

(34) Abcam. Ab65354 Superoxide Dismutase Activity Assay Kit (Colorimetric). 2019, 65354 (September), 19.

(35) Musachio, E. A. S.; Araujo, S. M.; Bortolotto, V. C.; de Freitas Couto, S.; Dahleh, M. M. M.; Poetini, M. R.; Jardim, E. F.; Meichtry, L. B.; Ramborger, B. P.; Roehrs, R.; Petri Guerra, G.; Prigol, M. Bisphenol A Exposure Is Involved in the Development of Parkinson like Disease in Drosophila Melanogaster. Food Chem. Toxicol. 2020, 137 (September 2019), 111128. 10.1016/j.fct.2020.111128.

(36) Information, P. Amplex ® Red Acetylcholine / Acetylcholinesterase Assay Kit (A12217). 2004, 8–11.

(37) Chen, M.; Hao, Y.; Chen, S. A Protocol for Investigating Lipidomic Dysregulation and Discovering Lipid Biomarkers from Human Serums. STAR Protoc. 2022, 3 (1), 101125. 10.1016/j.xpro.2022.101125.

(38) Chappel, J. R.; Kirkwood-Donelson, K. I.; Reif, D. M.; Baker, E. S. From Big Data to Big Insights: Statistical and Bioinformatic Approaches for Exploring the Lipidome. Anal. Bioanal. Chem. 2024, 416 (9), 2189–2202. 10.1007/s00216-023-04991-2.

(39) Aoyagi, R.; Ikeda, K.; Isobe, Y.; Arita, M. Comprehensive Analyses of Oxidized Phospholipids Using a Measured MS/MS Spectra Library. J. Lipid Res. 2017, 58 (11), 2229–2237. 10.1194/jlr.D077123.

(40) Esterbauer, H.; Cheeseman, K. H.; Dianzani, M. U.; Poli, G.; Slater, T. F. Separation and Characterization of the Aldehydic Products of Lipid Peroxidation Stimulated by ADP-Fe2+ in Rat Liver Microsomes. Biochem. J. 1982, 208 (1), 129–140. 10.1042/bj2080129.

(41) Pérez-Estrada, J. R.; Hernández-García, D.; Leyva-Castro, F.; Ramos-León, J.; Cuevas-Benítez, O.; Díaz-Muñoz, M.; Castro-Obregón, S.; Ramírez-Solís, R.; García, C.; Covarrubias, L. Reduced Lifespan of Mice Lacking Catalase Correlates with Altered Lipid Metabolism without Oxidative Damage or Premature Aging. Free Radic. Biol. Med. 2019, 135 (October 2018), 102–115. 10.1016/j.freeradbiomed.2019.02.016.

(42) Zheng, M.; Liu, Y.; Zhang, G.; Yang, Z.; Xu, W.; Chen, Q. The Applications and Mechanisms of Superoxide Dismutase in Medicine, Food, and Cosmetics. Antioxidants (Basel, Switzerland) 2023, 12 (9). 10.3390/antiox12091675.

(43) Li, W.; Pan, X.; Li, M.; ling, L.; Zhang, M. M.; liu, Z.; Zhang, K.; Guo, J.; Wang, H. Impact of Age on the Rotenone-Induced Sporadic PD Model Using Drosophila Melanogaster. Neurosci. Lett. 2023, 805 (March), 137187. 10.1016/j.neulet.2023.137187.

(44) Coulom, H.; Birman, S. Chronic Exposure to Rotenone Models Sporadic PD in Drosophila Melanogaster. J. Neurosci. Off. J. Soc. Neurosci. 2004, 24 (48), 10993– 10998. 10.1523/JNEUROSCI.2993-04.2004.

(45) Carpinella, I.; Crenna, P.; Calabrese, E.; Rabuffetti, M.; Mazzoleni, P.; Nemni, R.; Ferrarin, M. Locomotor Function in the Early Stage of PD. IEEE Trans. neural Syst. Rehabil. Eng. a Publ. IEEE Eng. Med. Biol. Soc. 2007, 15 (4), 543–551. 10.1109/TNSRE.2007.908933.

(46) van Hilten, J. J.; Hoogland, G.; van der Velde, E. A.; Middelkoop, H. A.; Kerkhof, G. A.; Roos, R. A. Diurnal Effects of Motor Activity and Fatigue in PD. J. Neurol. Neurosurg. Psychiatry 1993, 56 (8), 874–877. 10.1136/jnnp.56.8.874.

(47) Tong, T.; Duan, W.; Xu, Y.; Hong, H.; Xu, J.; Fu, G.; Wang, X.; Yang, L.; Deng, P.; Zhang, J.; He, H.; Mao, G.; Lu, Y.; Lin, X.; Yu, Z.; Pi, H.; Cheng, Y.; Xu, S.; Zhou, Z. Paraquat Exposure Induces Parkinsonism by Altering Lipid Profile and Evoking Neuroinflammation in the Midbrain. Environ. Int. 2022, 169, 107512. 10.1016/j.envint.2022.107512.

(48) Xicoy, H.; Brouwers, J. F.; Wieringa, B.; Martens, G. J. M. Explorative Combined Lipid and Transcriptomic Profiling of Substantia Nigra and Putamen in PD. Cells 2020, 9 (9). 10.3390/cells9091966.

(49) Wu, Y.; Wang, J.; Deng, Y.; Angelov, B.; Fujino, T.; Hossain, M. S.; Angelova, A. Lipid and Transcriptional Regulation in a PD Mouse Model by Intranasal Vesicular and Hexosomal Plasmalogen-Based Nanomedicines. Adv. Healthc. Mater. 2024, 13 (14), 1–14. 10.1002/adhm.202304588.

(50) Kalecký, K.; Bottiglieri, T. Targeted Metabolomic Analysis in PD Brain Frontal Cortex and Putamen with Relation to Cognitive Impairment. *npj Park*. Dis. 2023, 9 (1). 10.1038/s41531-023-00531-y.

(51) Raghu, P.; Joseph, A.; Krishnan, H.; Singh, P.; Saha, S. Phosphoinositides: Regulators of Nervous System Function in Health and Disease. Front. Mol. Neurosci. 2019, 12, 208. 10.3389/fnmol.2019.00208.

(52) Falabella, M.; Vernon, H. J.; Hanna, M. G.; Claypool, S. M.; Pitceathly, R. D. S. Cardiolipin, Mitochondria, and Neurological Disease. Trends Endocrinol. Metab. 2021, 32 (4), 224–237. 10.1016/j.tem.2021.01.006.

(53) Paradies, G.; Paradies, V.; Ruggiero, F. M.; Petrosillo, G. Role of Cardiolipin in Mitochondrial Function and Dynamics in Health and Disease: Molecular and Pharmacological Aspects. Cells 2019, 8 (7). 10.3390/cells8070728.

(54) Pokhrel, S. No TitleΕΛΕΝΗ. Αγαη 2024, 15 (1), 37–48.

(55) Sanz Muñoz, S.; Marlet, F.; Bilgin, M.; Dreier, J.; Bezard, E.; Dehay, B.; Jaunmuktane, Z.; Maeda, K.; Galvagnion, C. Lipidomic Profiling Reveals Shared and Distinct Pathological Signatures in Sporadic PD and GBA Mutation Carriers: Implications for Disease Mechanisms; 2024. 10.1101/2024.10.11.617800.

(56) Worth, A. J.; Basu, S. S.; Snyder, N. W.; Mesaros, C.; Blair, I. A. Inhibition of Neuronal Cell Mitochondrial Complex i with Rotenone Increases Lipid β-Oxidation, Supporting Acetyl-Coenzyme a Levels. J. Biol. Chem. 2014, 289 (39), 26895–26903. 10.1074/jbc.M114.591354.

(57) Al-Zaid, F. S.; Hurley, M. J.; Dexter, D. T.; Gillies, G. E. Neuroprotective Role for RORA in PD Revealed by Analysis of Post-Mortem Brain and a Dopaminergic Cell Line. NPJ Park. Dis. 2023, 9 (1), 119. 10.1038/s41531-023-00563-4.

(58) Fernández-Irigoyen, J.; Cartas-Cejudo, P.; Iruarrizaga-Lejarreta, M.; Santamaría, E. Alteration in the Cerebrospinal Fluid Lipidome in PD: A Post-Mortem Pilot Study. Biomedicines 2021, 9 (5). 10.3390/biomedicines9050491.

(59) Yakunin, E.; Kisos, H.; Kulik, W.; Grigoletto, J.; Wanders, R. J. A.; Sharon, R. The Regulation of Catalase Activity by PPAR γ Is Affected by α-Synuclein. Ann. Clin. Transl. Neurol. 2014, 1 (3), 145–159. 10.1002/acn3.38.

(60) Djordjević, V. V; Kostić, J.; Krivokapić, Ž.; Krtinić, D.; Ranković, M.; Petković, M.; Ćosić, V. Decreased Activity of Erythrocyte Catalase and 1196 Glutathione Peroxidase in Patients with Schizophrenia. Medicina (Kaunas). 2022, 58 (10). 10.3390/medicina58101491.

(61) Poleto, K. H.; Janner, D. E.; Dahleh, M. M. M.; Poetini, M. R.; Fernandes, E. J.; Musachio, E. A. S.; de Almeida, F. P.; Amador, E. C. de M.; Reginaldo, J. C.; Carriço, M. R. S.; Roehrs, R.; Prigol, M.; Guerra, G. P. P-Coumaric Acid Potential in Restoring Neuromotor Function and Oxidative Balance through the Parkin Pathway in a Parkinson Disease-like Model in Drosophila Melanogaster. Food Chem. Toxicol. an Int. J. Publ. Br. Ind. Biol. Res. Assoc. 2024, 115002. 10.1016/j.fct.2024.115002.

(62) Kawahata, I.; Yagishita, S.; Kazuko, H.; Nagatsu, I.; Nagatsu, T.; Ichinose, H. Immunohistochemical Analyses of the Postmortem Human Brains from Patients with PD with Anti-Tyrosine Hydroxylase Antibodies; 2015.

(63) Dexter, D. T.; Carter, C. J.; Wells, F. R.; Javoy□Agid, F.; Agid, Y.; Lees, A.; Jenner, P.; Marsden, C. D. Basal Lipid Peroxidation in Substantia Nigra Is Increased in PD. J. Neurochem. 1989, 52 (2), 381–389. 10.1111/j.1471-4159.1989.tb09133.x.

(64) Yang, W.; Chang, Z.; Que, R.; Weng, G.; Deng, B.; Wang, T.; Huang, Z.; Xie, F.; Wei, X.; Yang, Q.; Li, M.; Ma, K.; Zhou, F.; Tang, B.; Mok, V. C. T.; Zhu, S.; Wang, Q. Contra-Directional Expression of Plasma Superoxide Dismutase with Lipoprotein Cholesterol and High-Sensitivity C-Reactive Protein as Important Markers of PD Severity. Front. Aging Neurosci. 2020, 12, 53.

(65) Carrillo, F.; Palomba, N. P.; Ghirimoldi, M.; Didò, C.; Fortunato, G.; Khoso, S.; Giloni, T.; Santilli, M.; Bocci, T.; Priori, A.; Pietracupa, S.; Modugno, N.; Barberis, E.; Manfredi, M.; Signorelli, P.; Esposito, T. Multiomics Approach Discloses Lipids and Metabolites Profiles Associated to PD Stages and Applied Therapies. Neurobiol. Dis. 2024, 202, 106698.

(66) Galper, J.; Dean, N. J.; Pickford, R.; Lewis, S. J. G.; Halliday, G. M.; Kim, W. S.; Dzamko, N. Lipid Pathway Dysfunction Is Prevalent in Patients with PD. Brain 2022, 145 (10), 3472–3487. 10.1093/brain/awac176.

(67) Hirano, S.; Shinotoh, H.; Aotsuka, A.; Sato, K.; Tanaka, N.; Ota, T.; Asahina, M.; Fukushi, K.; Kuwabara, S.; Hattori, T.; Suhara, T.; Irie, T. Mapping of Brain Acetylcholinesterase Alterations in Lewy Body Disease by PET. Neurology 2009, 73 (4), 273–278. 10.1212/WNL.0b013e3181ab2b58.

