## Supplementary information for "Exposure to rotenone triggers redox driven system-wide lipidome alterations and metabolic trade-offs linked to Parkinson’s disease"

**Figures S1 to S9**

**Exposure to rotenone triggers redox driven system-wide lipidome alterations and metabolic trade-offs linked to Parkinson’s disease**

Ashutosh K. Tiwari^a#^, Priya Rathor^a,b#^, Rajendra Patel^c^, Pawan Kumar Jha^d^, Nick Birse^e^, Ratnasekhar CH^a,b,e*^

^a^ Metabolomics laboratory, CSIR-Central institute of Medicinal & Aromatic Plants (CIMAP), Lucknow-226015, India

^b^ Academy of Scientific and Innovative research, Delhi- 201 002, India

^c^ Biological central laboratory, CSIR-CIMAP, Lucknow-226015, India

^d^ Department of Systems Pharmacology & Translational Therapeutics, Perelman School of Medicine, University of Pennsylvania, Philadelphia, PA, 19104, USA

^e^School of biological Sciences laboratory, Queen’s University Belfast, BT9 5DL, United Kingdom

^#^ These authors contributed equally

**^*^ *Correspondence*:**  Dr. Ratnasekhar Ch, PhD

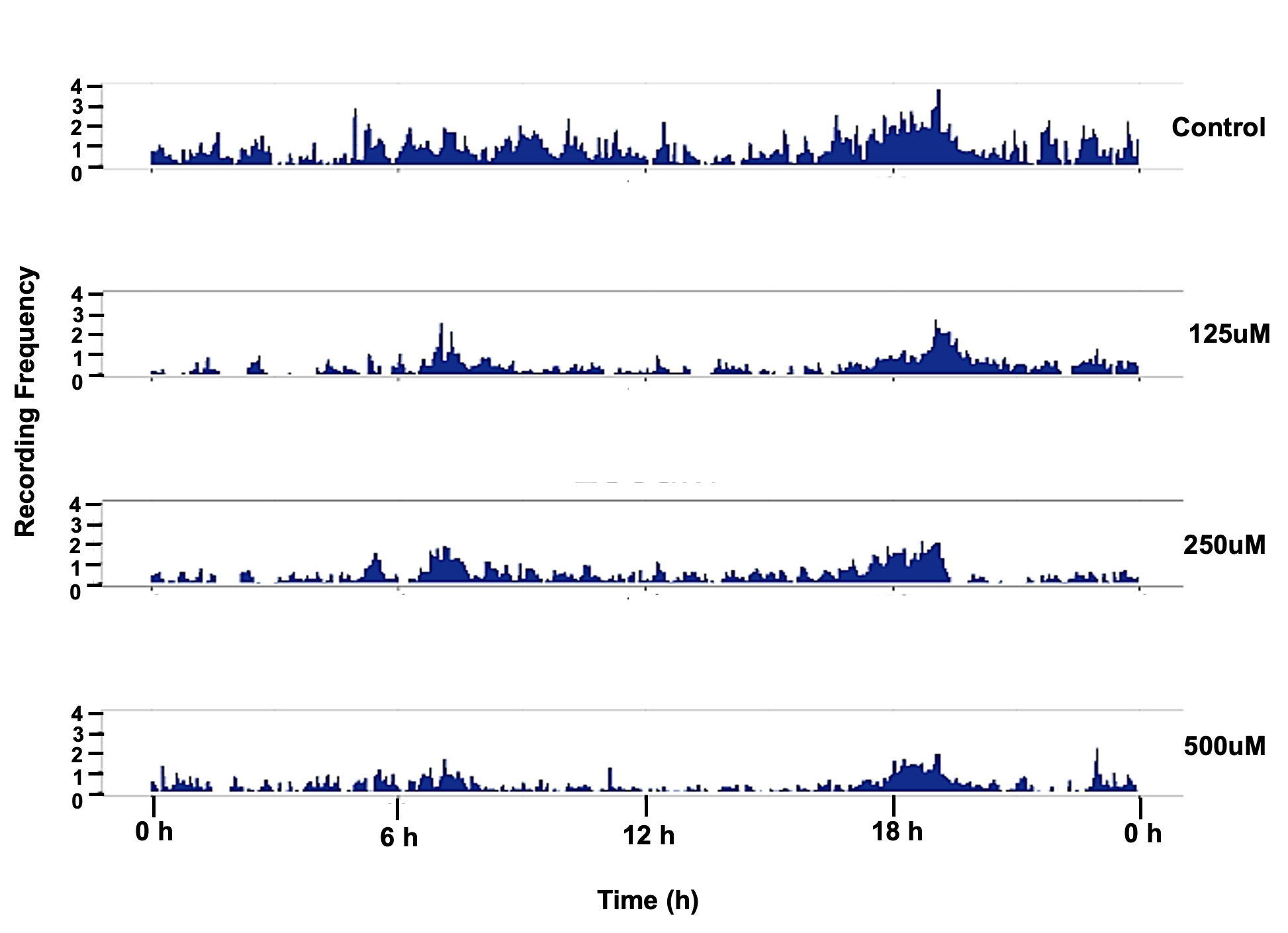

Figure S1: Typical actograms represent the biological variables under chronic exposure of

rotenone in 12:12-hour LD cycles. Each actogram of rotenone (125uM, 250uM and 500uM)

exposed flies represent an activity profile during 24 hours.

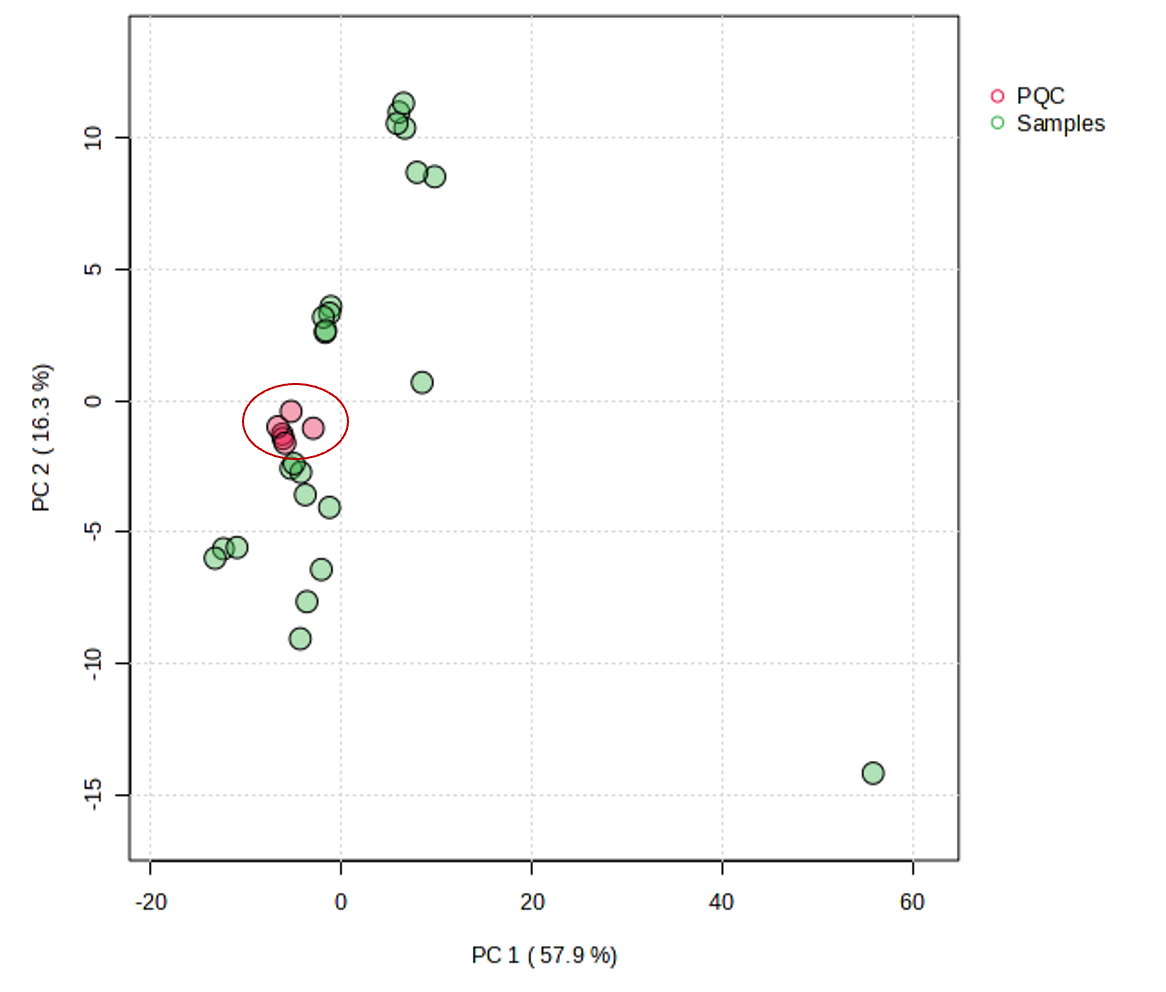

Figure S2: PCA score plot of samples and pooled QC.

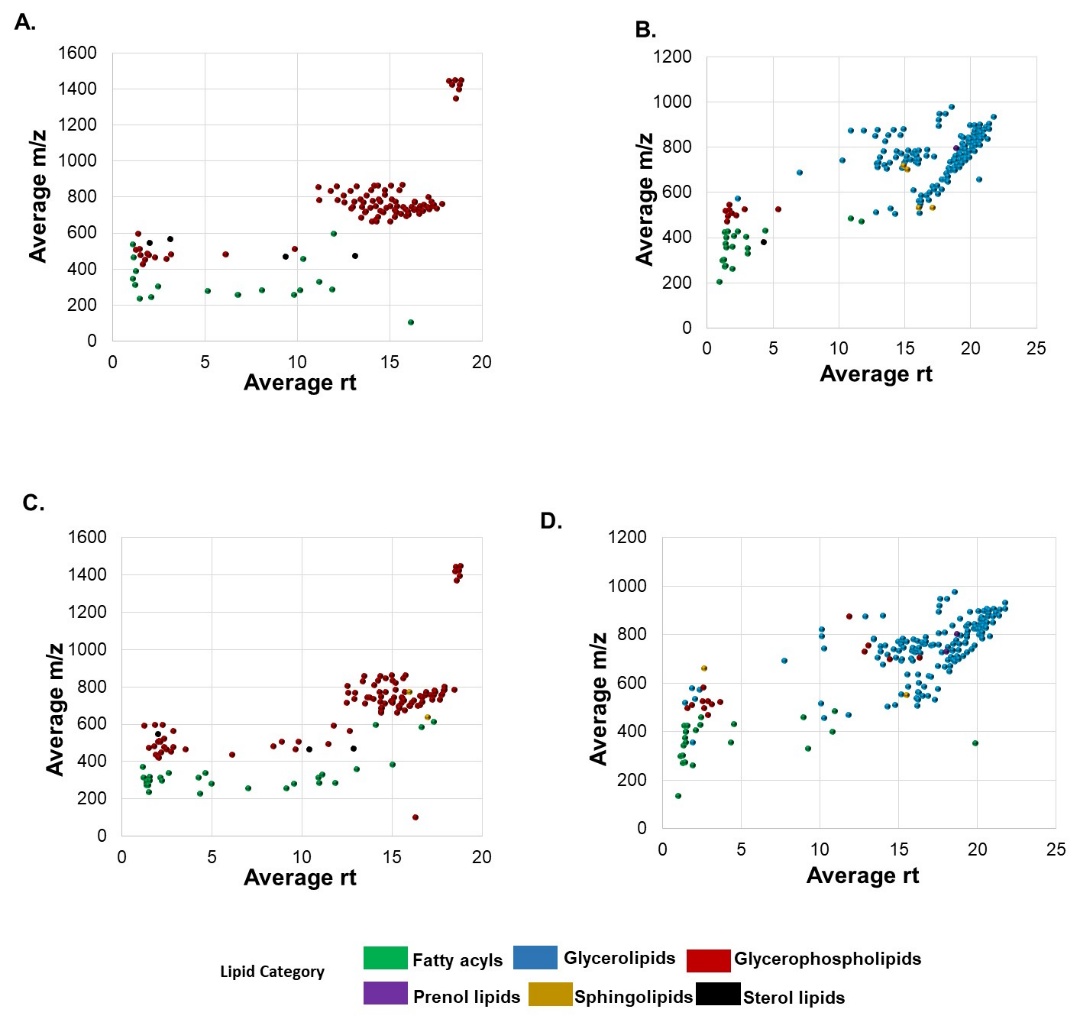

Figure S3: Scatter plot showing the m/z and average rt of detected lipids in brain and

body tissue A. ESI Negative of brain tissue B. ESI Positive of brain tissue C. ESI Negative of

body tissue D. ESI Positive of body tissue

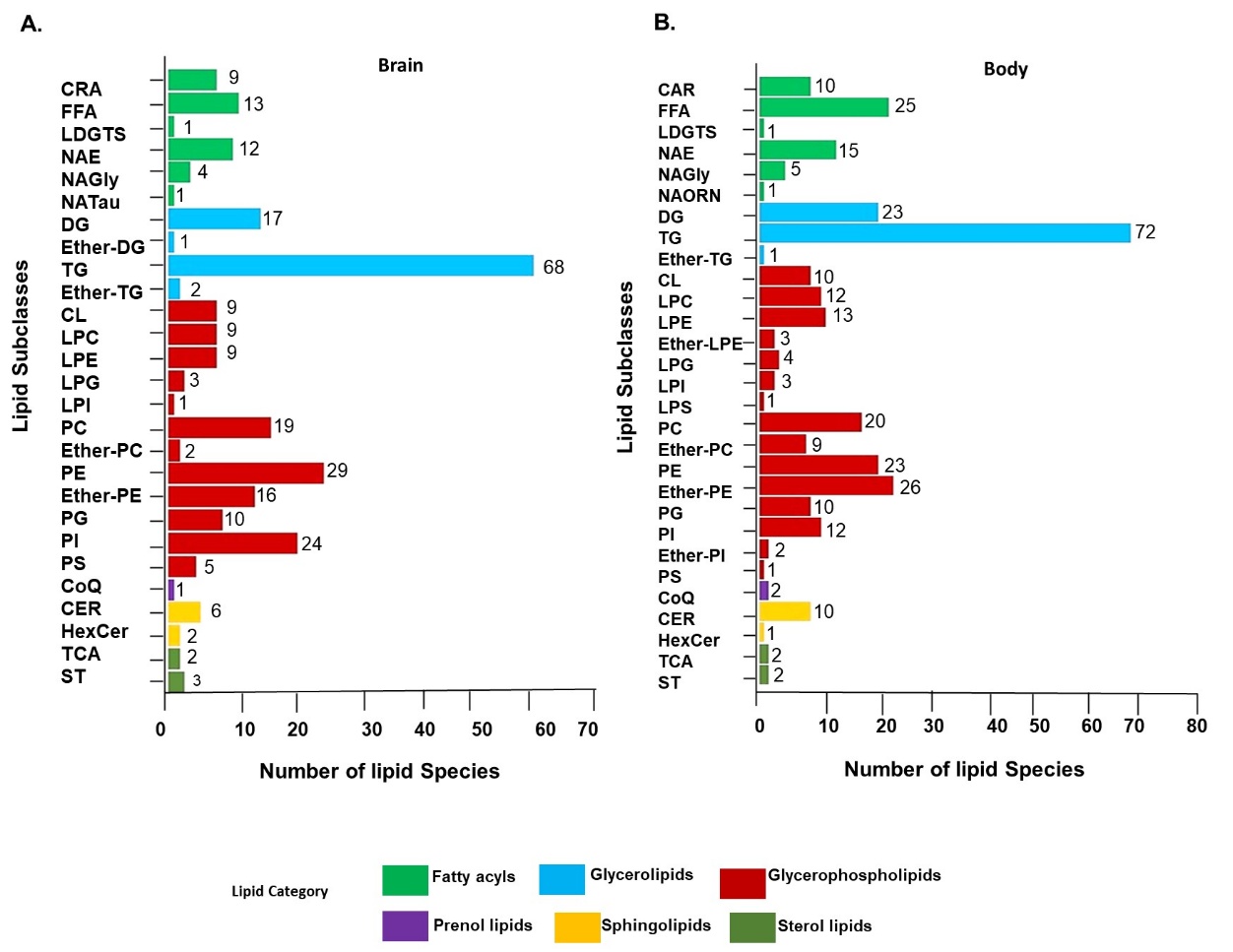

**Figure S4**: Statistical plot for lipid subclasses in brain and body tissues of D. melanogaster

exposed with rotenone. A. 27 subclass of lipids existing in brain tissue and B. 29 subclass of lipids existing in body tissue. CAR: Acylcarnitine, FFA: Free fatty acids, LDGTS: Lysodiacylglyceryltrimethylhomoserine, NAE: N-acyl ethanolamines, NAGly: N-acyl glycyl serine, NATau: N-acyl taurine, NAOrn: N-acyl ornithine, DG: Diacylglycerol, Ether-DG: Ether-linked diacylglycerol, TG: Triacylglycerol, Ether-TG: Ether-linked triacylglycerol, CL: Cardiolipin, LPC: Lysophosphatidylcholine, LPE: Lysophosphatidylethanolamine, LPG:  Lysophosphatidylglycerol, LPI: Lysophosphatidylinositol, PC: Phosphatidylcholine, Ether-PC: Ether-linked phosphatidylcholine, PE: Phosphatidylethanolamine, Ether-PE: Ether-linked phosphatidylethanolamine, PG: Phosphatidylglycerol, PI: Phosphatidylinositol, PS: Phosphatidylserine, CoQ: Coenzyme Q, CER: Ceramides, HexCer: Hexosylceramides, TCA: Taurocholic acid, ST: Sterol derived lipids, Ether-PI: Ether-linked Phosphatidylinositol, Ether-LPE: Ether-linkedlysophosphatidylethanolamine.

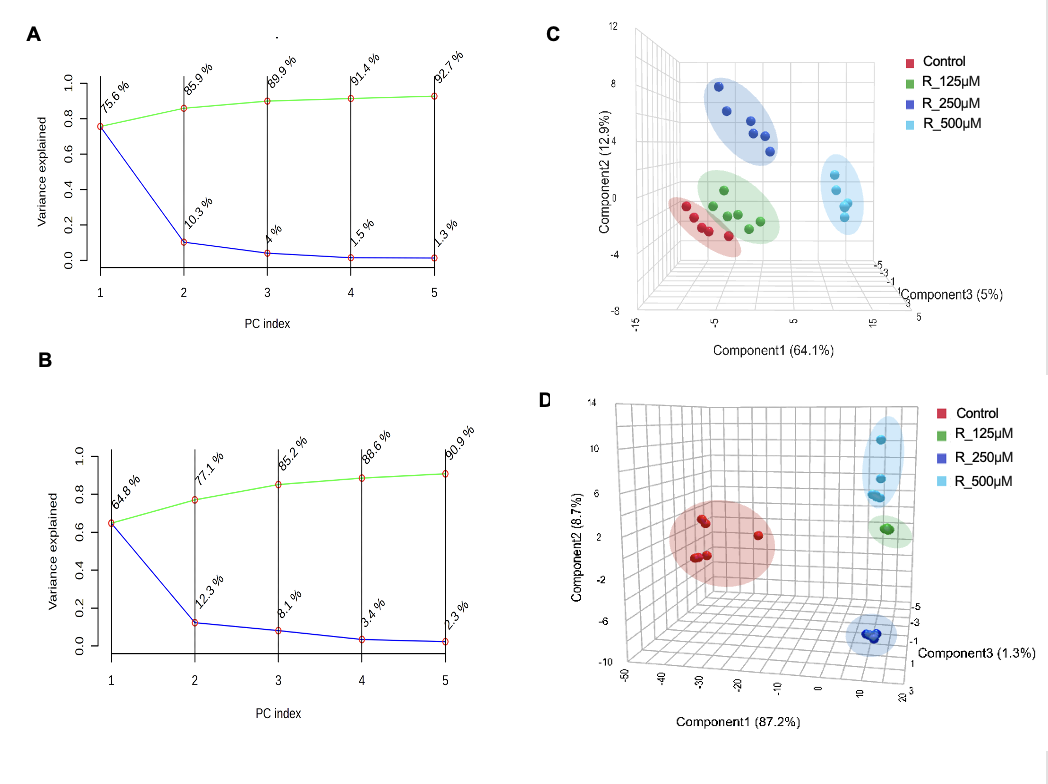

Figure S5: Individual and cumulative explained variances of principal components for A) Drosophila brain samples in ESI (-) mode. B) Drosophila brain samples in ESI (+) mode. The cumulative (green line) explained variance shows the accumulation of variance for each principal component number. The individual (blue line) explained variance describes the variance of reach principal component. C) PLS-DA analysis of control and rotenone treated brain tissue samples in ESI (-) mode. D) PLS-DA analysis of control and rotenone treated brain tissue samples in ESI (+) mode.

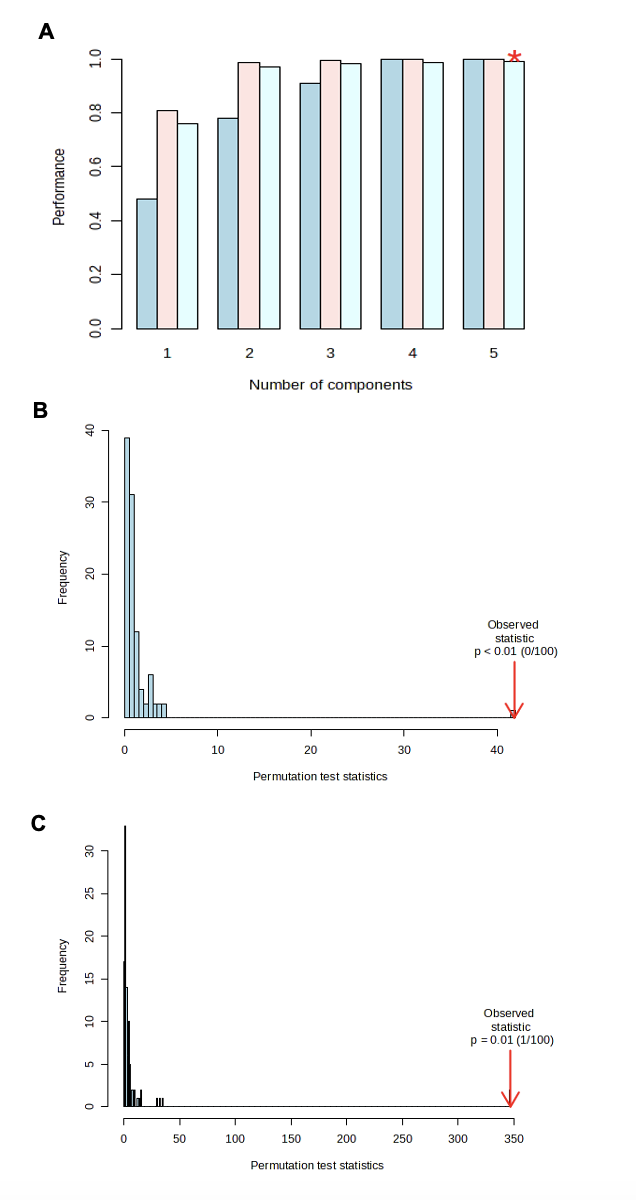

Figure S6: A) Multivariate analysis using PLS-DA cross validation. Bar plots showing the three performance measures (prediction accuracy, R2 and Q2) using different number of components. The red ‘*’ indicates the best values of the currently selected measures (Q2). (B) Statistical validation of the PLS-DA by permutation analysis using 1000 different model permutations in ESI (-) mode for brain tissue samples. C) (B) Statistical validation of the PLS-DA by permutation analysis using 1000 different model permutations in ESI (+) mode for brain tissue samples. The goodness of fit and predictive capability of the original class assignments is much higher compared to ratios based on the permutation class assignments (P < 0.001).

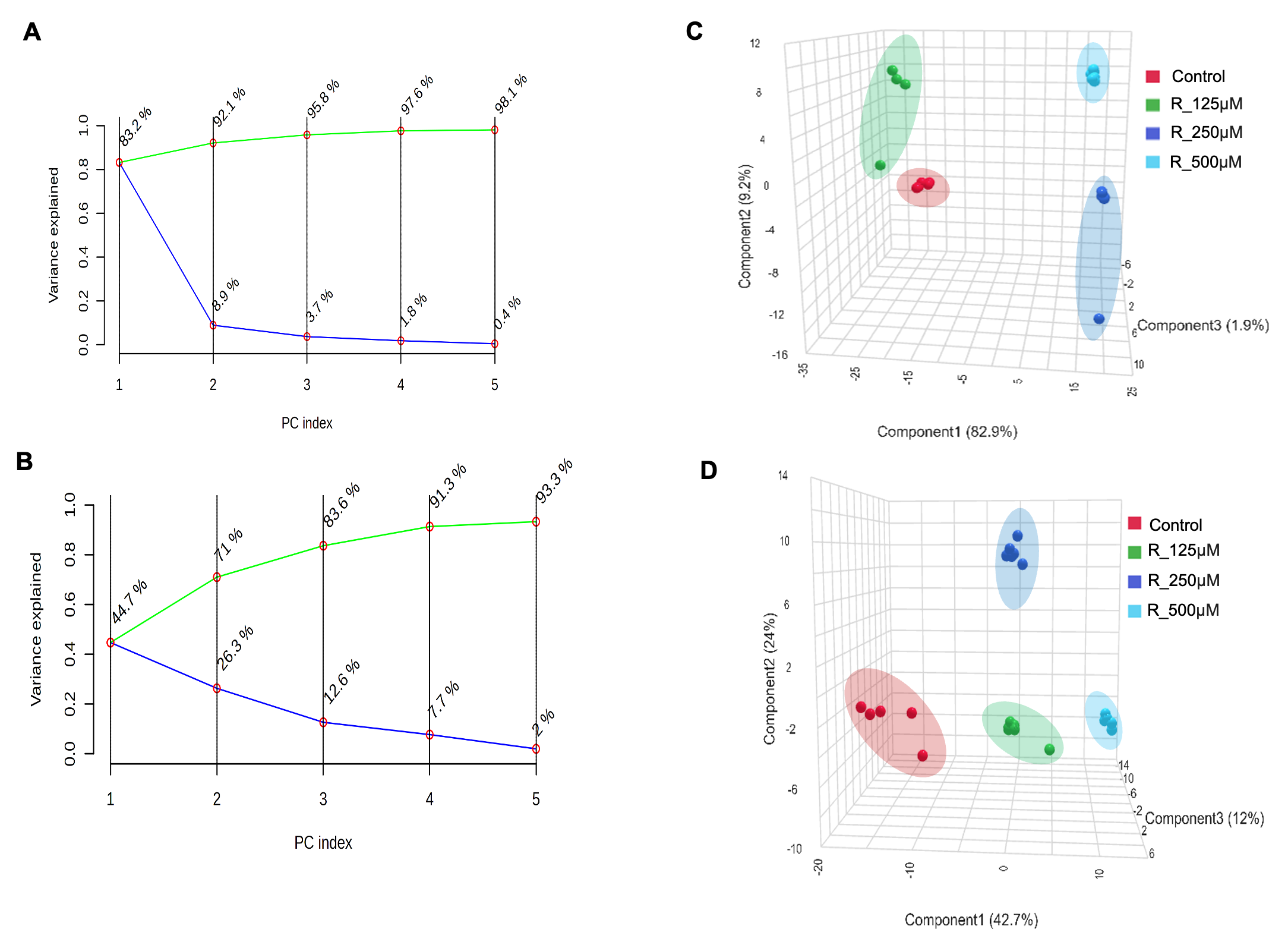

Figure S7: Individual and cumulative explained variances of principal components for A) *Drosophila* body samples in ESI (-) mode. B) Drosophila body samples in ESI (+) mode. The cumulative (green line) explained variance shows the accumulation of variance for each principal component number. The individual (blue line) explained variance describes the variance of reach principal component. C) PLS-DA analysis of control and rotenone treated body tissue samples in ESI (-) mode. D) PLS-DA analysis of control and rotenone treated body tissue samples in ESI (+) mode.

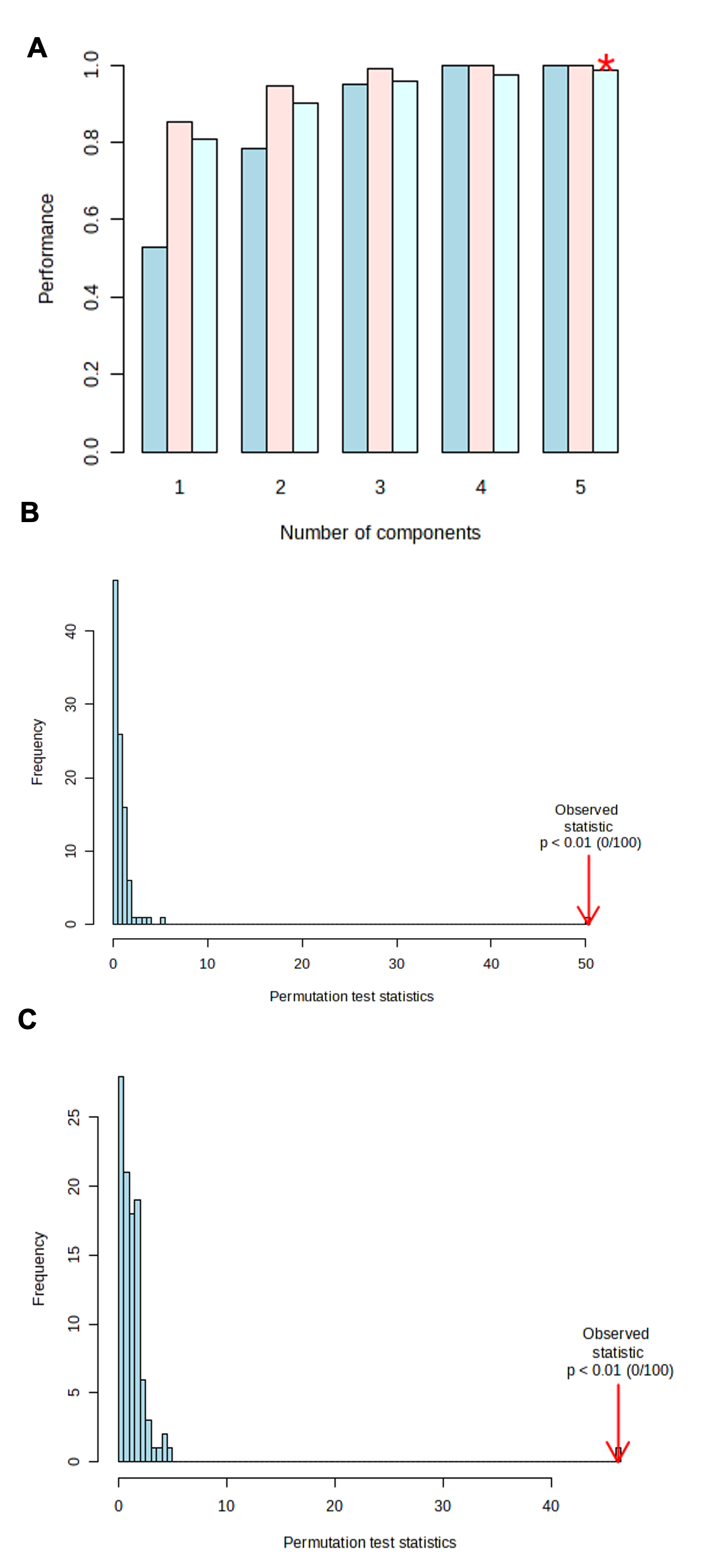

Figure S8: A) Multivariate analysis using PLS-DA cross validation. Bar plots showing the three performance measures (prediction accuracy, R2 and Q2) using different number of components. The red ‘*’ indicates the best values of the currently selected measures (Q2). (B) Statistical validation of the PLS-DA by permutation analysis using 1000 different model permutations in ESI (-) mode for body tissue samples. C) (B) Statistical validation of the PLS-DA by permutation analysis using 1000 different model permutations in ESI (+) mode for body tissue samples. The goodness of fit and predictive capability of the original class assignments is much higher compared to ratios based on the permutation class assignments (P < 0.001).

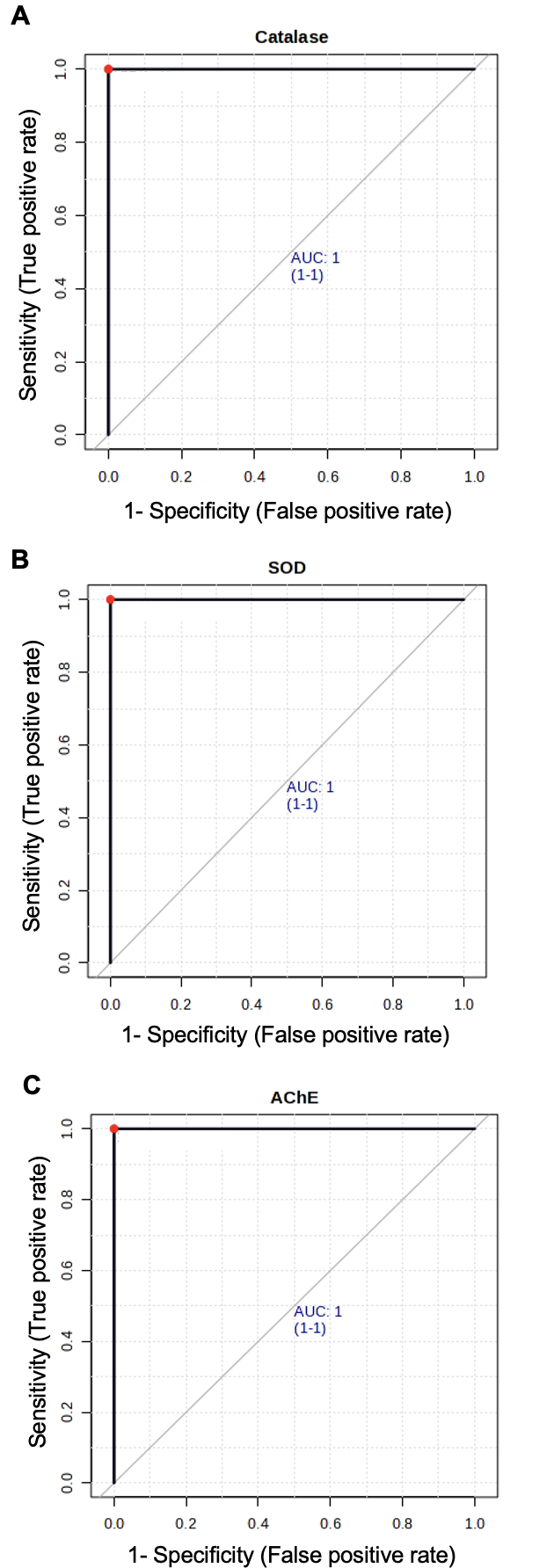

**Figure S9:** Receiving operating characteristic curve performance of rotenone exposed *Drosophila* samples. A) Catalase B) SOD C) AChE.
